# Avian Diversity and Community Structure in the Peri-urban Landscape of Magadh University, Bodh Gaya

**DOI:** 10.64898/2026.09.17.752356

**Authors:** Manisha Kumari, Partha Pratim Das, Kumari Aditi

**Author notes:** Corresponding Authors: **Dr. Kumari Aditi**, Senior Assistant Professor, Post Graduate Department of Zoology, Magadh University, Bodh Gaya, India – 824234, Contact no, **Dr. Partha Pratim Das**, Senior Assistant Professor, Post Graduate Department of Chemistry, Magadh University, Bodh Gaya, India – 824234.

## Abstract

University campuses function as important refugia for biodiversity within an increasingly urbanizing landscape. Magadh University (MU) is among the oldest universities of Bihar, India, located in Bodh Gaya, a place of spiritual tourism, near a UNESCO heritage site. The campus hosts a mosaic of heterogeneous habitats, rich vegetation and dense canopy cover along with varied range of anthropogenic interferences. This study mainly focusses on measuring the alpha diversity of avifaunal species and persistence across four seasons from July 2025 to June 2026, in the peri-urban landscape of MU. A total of 78 bird species belonging to 41 families and 16 orders were recorded using a combination of point count, line transect and acoustic monitoring. The study revealed that the order Passeriformes was the most species-rich (47.44%). Muscicapidae was the most species-rich family (6 species). Relatively high Shannon diversity (H’) = 3.278, low Simpson’s dominance (D = 0.062) and a high Pielou’s evenness (J) = 0.751 indicated the avian community was distributed across diverse assemblage. Species richness estimates from Chao1 (78.5), iChao1 (79.02) and ACE (79.53) were comparable to recorded 78 species, indicating sampling completeness. Observed species were further classified into five major feeding guilds, majority of which were insectivorous species. 77 of the observed species were classified as Least Concern while *Coracias benghalensis* was categorized as Near Threatened. These findings demonstrate that the MU campus supports significant taxonomic, functional and temporal avian assemblage, highlighting the conservation values of the campus landscapes within rapidly urbanizing regions.

## Introduction

Birds constitute an important component of terrestrial vertebrate communities and contribute to a broad range of ecological processes that influence ecosystem functioning and stability (Whelan et al. 2015). Variation in their morphology, behaviour, habitat use and feeding strategies enables different species to occupy distinct ecological niches and perform diverse ecological functions (Gaston 2022; Pena et al. 2022; Diaz-Chaux et al. 2025; Chen et al. 2026). Birds are also widely used as indicators of environmental conditions, as changes in their composition and abundance can reflect variations in habitat quality, resource availability and other environmental factors (Egwumah et al. 2017; Mekonen 2017). Certain species have additionally been recognized as flagship, keystone, umbrella or focal taxa, making their conservation relevant to other associated components of biological communities (Padoa-Schioppa et al. 2006; Fraixedas et al. 2020; Wang et al. 2023).

The ecological functions performed by birds include seed and pollen dispersal, scavenging, predation of insects and small vertebrates, weed control, nutrient cycling and facilitation of vegetation regeneration and ecological restoration (Gaston 2022). These interactions can influence vegetation dynamics, agricultural productivity and ecosystem recovery (Whelan et al. 2008; Gaston 2022). Birds also have socioeconomic and cultural value through activities such as birdwatching, wildlife photography, ecotourism and environmental education (Schwoerer and Dawson 2022; Larasati et al. 2024; Saha and Chakraborty 2025). However, these ecological and social values are increasingly affected by habitat loss and modification. Urban expansion, deforestation and habitat fragmentation have reduced natural habitats and altered the availability of resources required by many bird species (McKinney 2002). While some adaptable species persist in synanthropic environments, species with more specialized habitat requirements may be more vulnerable to local population declines. At the same time, heterogeneous urban habitats containing mature trees, wetlands, grasslands and shrub-dominated areas can retain resources and structural features that support birds and other wildlife (Guthula et al. 2022; Iwachido et al. 2023).

University campuses represent a type of urban landscape where biodiversity can persist alongside intensive human use (Colding and Barthel 2017; Liu et al. 2021; Sanllorente et al. 2023; Kirazli et al. 2025). Many campuses contain plantations, gardens, open grounds, wetlands and other vegetated areas that provide feeding, nesting, roosting and breeding opportunities for birds (Devi et al. 2012; Rajendran et al. 2014; Rathod and Bhaduri 2022; Shivhare et al. 2022; Kumar et al. 2024; Singh et al. 2024; Imran et al. 2025; Kamboj et al. 2026). Their relatively stable land-use boundaries and accessibility also make them suitable locations for repeated biodiversity surveys and long-term ecological monitoring (Zhang et al. 2018; Guthula et al. 2022). Records of diverse avian communities from university campuses suggest that these areas can contribute to the retention of biodiversity within landscapes increasingly dominated by urban or agricultural land use (Guthula et al. 2022).

The conservation value of such campuses is likely to be high where mature vegetation and heterogeneous habitat is retained. Trees can provide nesting and roosting sites, foraging substrates and arboreal microhabitats, while ground vegetation, shrubs and water bodies add further habitat components. The combination of these features can support resident and seasonally occurring bird species. University campuses have consequently been discussed in relation to Other Effective Area-Based Conservation Measures (OECMs), particularly where their management contributes to the *in situ* conservation of biodiversity (Saraniya et al. 2026). However, the conservation significance of an individual campus needs to be considered on the basis of site-specific information on species composition, abundance, habitat characteristics, occurrence patterns and persistence.

Within this broader context, the MU campus at Bodh Gaya provides an appropriate setting for examining avian diversity within a densely populated but vegetation-rich landscape. India supports a highly diverse avifauna accounting for nearly 13% of global avian diversity, and there are approximately 1,389 recorded bird species (Callaghan et al. 2021, eBird 2026). Bihar also supports considerable avian diversity, with about 483 species reported from the state and approximately 170 species documented from Gaya Ji district (Maheswaran and Alam 2025; eBird 2026). The geographical location of Bodh Gaya further adds ecological relevance because the region occurs within a landscape associated with several natural and semi-natural ecosystems, including the Brahmayoni Hills and the Falgu River. The surrounding habitats such as Rajgir Hills, Kaimur-Rohtas hill range and Gautam Buddha Wildlife Sanctuary may provide additional resources and movement pathways for birds at the local and regional scales (Rodgers 2000).

Bodh Gaya is internationally recognized for its rich cultural and historical legacy, anchored by the Mahabodhi Temple Complex, a UNESCO World Heritage Site (UNESCO 2002). The interaction of cultural landscapes, urban development, institutional areas and surrounding natural habitats provides an opportunity to examine the persistence of biodiversity within a landscape subject to multiple forms of human use. Despite this ecological setting, information on the composition and structure of avian community within the peri-urban MU campus remains limited. A systematic assessment of the campus avifauna is therefore relevant for documenting its bird community and examining the extent to which different campus habitats are used by birds.

The MU campus has retained extensive tree cover and variety of vegetated habitats over several decades, and has undergone considerably less habitat modification than the surrounding urban matrix. The campus contains mature trees, shrublands, grasslands, landscaped gardens, orchards, wetlands, ponds, and built-up areas, forming a heterogeneous habitat mosaic. Such structural variation provides different resources and microhabitats for birds with contrasting ecological requirements. The present study therefore aimed to characterize the avian community of the MU campus through year-round field observations, with particular emphasis on the alpha diversity, relative abundance, residential status, habitat associations, feeding guild composition, and temporal patterns of occurrence, within a peri-urban landscape. We also examined the potential contribution of MU campus in local avifaunal conservation. Our findings provide a baseline for future biodiversity assessments and may highlight specific habitat variables that help to prioritize for targeted habitat preservation and enhancement.

## Materials and methods

### Study Area

This study was carried out at MU campus, situated in the Bodh Gaya region of Gaya Ji district, Bihar, India (Fig. 1a-c). The campus covers approximately 303.17 acres (122.7 ha) and extends between 24.40-24.41°N and 84.57°E, with a mean elevation of 149.48 m above sea level (Fig. 1c). Gaya-Patna National Highway-83 (NH-83) borders the campus on its western side, whereas agricultural land occurs predominantly along its eastern boundary.

**Fig. 1.**
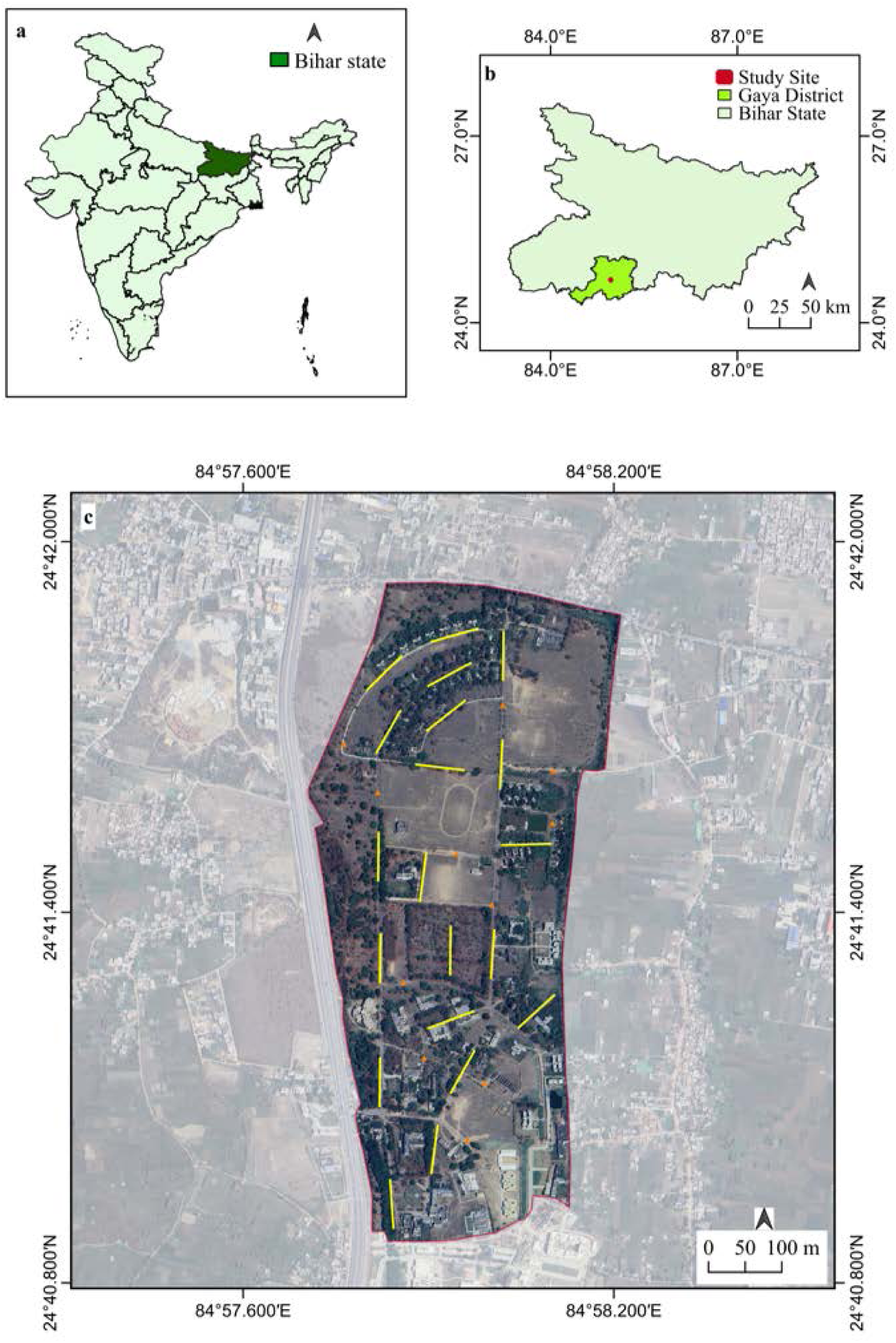
Location map of the study area. (a) Geographic location of Bihar state within India. (b) Location of Gaya Ji district within Bihar, with the Magadh University (MU) campus highlighted in red (c) Satellite image of the MU campus, Bodh Gaya, showing the study area outlined in red. 150m long line transects represented by yellow lines and 11 stationary point count stations represented by orange triangle were fixed for the study. Geographic coordinates, scale bars and north arrows are provided for spatial reference.

The campus contains a combination of vegetated and built-up areas, with considerable variation in vegetation structure and land use. Survey sites were distributed across the major habitat types present within the campus, and their spatial distribution is presented in Fig. 2 (a-ab). These sites include mature trees, avenue plantations, woodland patches, gardens, grasslands, wetlands, shrub-dominated areas, open fields, sports grounds, vegetation around academic buildings, residential areas with mature trees, and peripheral woodland. The vegetation comprises dense cover of naturally established trees, scattered mature trees, shrubs, grass-covered areas, seasonal wetlands and open patches. This variation in habitat structure provided the principal setting for assessing bird occurrence and habitat use across the campus.

**Fig. 2.**
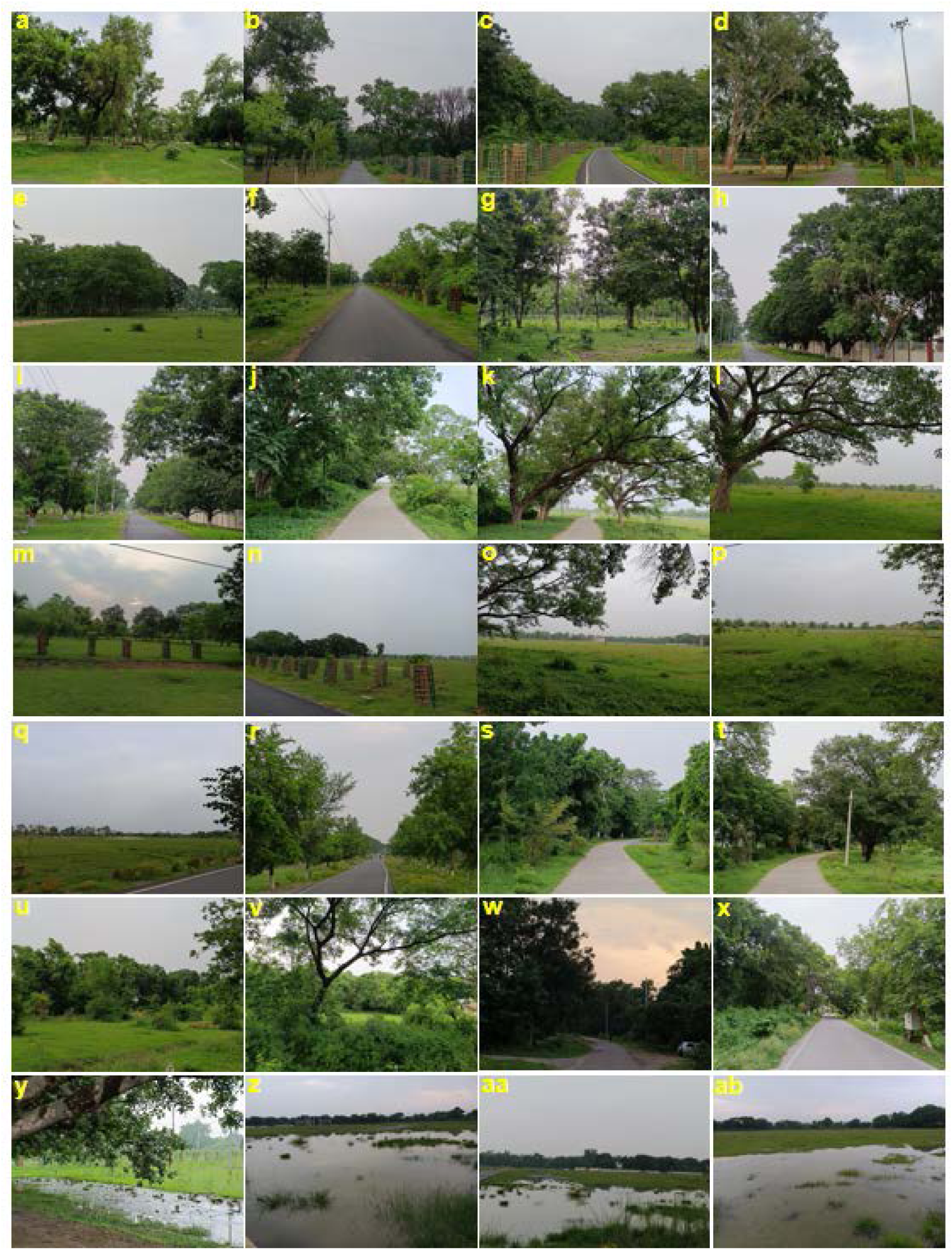
Representative sampling locations surveyed for avifaunal surveys across the MU campus. (a) Green spaces around academic buildings; (b, c) Avenue plantation in the Science Block; (d) Botanical garden; (e) *Terminalia arjuna* plantation adjoining open grassland; (f) Tree-lined campus roadway; (g) Dense mixed-species plantation of young trees; (h, i) Avenue plantation along the Guest House road; (j) Mixed tree cover adjacent to Sports field; (k) Mature *Albizia lebbeck* trees estimated to be over 50 years old; (l) Open grassland; (m) Grassland transitioning into peripheral woodland; (n-q) Sports field; (r) Avenue plantation bordering the road to the secondary entrance gate; (s-x) Mature tree cover surrounding residential areas; (y-ab) wetland areas. These representative habitats encompass the range of vegetation types and landscape features surveyed during the study and illustrate habitat heterogeneity that support diverse avian communities within the campus.

The campus vegetation comprises a diverse assemblage of trees, shrubs, herbs, grasses and aquatic or wetland-associated plants. Frequently occurring tree species include *Terminalia arjuna*, *Albizia lebbeck*, *Albizia saman*, *Azadirachta indica*, *Ficus benghalensis*, *Mangifera indica*, *Ficus religiosa*, *Ziziphus mauritiana*, *Syzygium cumini*, *Tamarindus indica*, *Pithecellobium dulce*, *Bauhinia variegata*, *Neolamarckia cadamba*, *Dalbergia sissoo*, *Lagerstroemia speciosa*, *Butea monosperma*, *Saraca asoca*, *Aegle marmelos, Bombax ceiba*, *Tectona grandis*, *Cassia fistula*, *Delonix regia*, *Peltophorum pterocarpum*, *Moringa oleifera*, *Vachellia nilotica*, *Prosopis juliflora*, *Phoenix sylvestris*, *Alstonia scholaris*, *Phyllanthus emblica*, *Putranjiva roxburghii*, *Swietenia macrophylla*, *Monoon longifolium*, *Morus alba, Eucalyptus* spp. and *Psidium guajava*. Other vegetation recorded inside the campus incudes *Bambusa* spp., *Cynodon dactylon*, *Lantana camara*, *Oplismenus burmanni*, *Stellaria media*, *Phyllanthus niruri*, *Phyllanthus urinaria*, *Rauvolfia tetraphylla*, *Cyperus* sp., *Parthenium* sp., and *Hyptis* spp. Plant identification was carried out with reference to Sahni (2000).

### Climatic Conditions

According to the Köppen climate classification, the study area experiences a humid subtropical (Cwa) climate characterized by hot, dry summers, cool winters, and rainfall concentrated during monsoon months. For the purpose of seasonal analyses, the annual climatic cycle was categorized into four seasons based on the prevailing local weather conditions: pre-monsoon (summer), monsoon, post-monsoon and winter. The pre-monsoon extends from March to June, with temperatures ranging from 25-46°C and cumulative precipitation of approximately 5-35 mm. The monsoon season occurs from July to September, recording temperatures between 25-35°C and receiving approximately 950-1100 mm of rainfall. The post-monsoon season spans October to November, with temperatures ranging from 18-30°C and precipitation of 40-77 mm. Winter extends from December to February and is characterized by temperatures between 3-25°C and precipitation ranging from 50-80 mm (Fig. 3a, b). Bird surveys were conducted across all four seasons to capture seasonal variation in avian diversity and community structure.

**Fig. 3.**
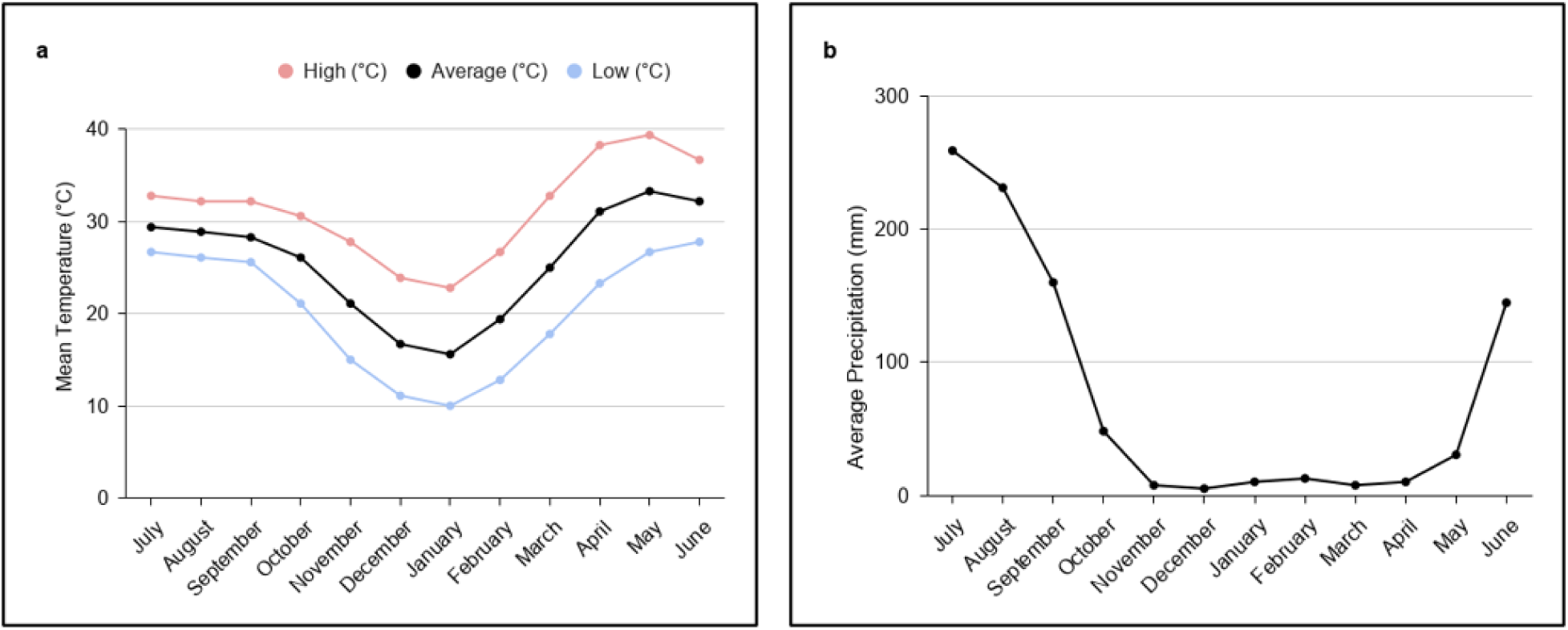
Monthly climatic profile of the study area. (a) Mean monthly maximum, average and minimum air temperature (°C) from July 2025 to June 2026. (b) Mean monthly precipitation (mm) from July 2025 to June 2026. The x-axis represents the months of the study period, whereas the y-axis indicates temperature (°C) in panel a and precipitation (mm) in panel b.

### Data collection

Avian surveys were conducted from June 2025 to May 2026 using a combination of point count, line transect and passive acoustic monitoring (PAM) methods following the established protocols (Javed and Kaul 2002; Rempel et al. 2005; Greenwood 2007). Point counts were conducted at 11 predetermined sites around the grassland and open fields with little canopy cover. Each of these sites were surveyed for 5 minutes, covering a visual radius of 50m. Line transect survey was conducted along 20 fixed tracks of 150m length, selected randomly throughout the campus such that majority of the study area was covered. Each of these transects were walked at a uniform pace. It was ensured that the individual birds were not counted twice. Passive acoustic recordings were obtained using Autonomous Recording Units (ARU; frequency range 50Hz to 12 kHz) deployed at an approximate height of 1.5 meters above ground level. Acoustic recordings were examined, and species identifications based on vocalizations were verified through direct field observations to minimize any false-positive detections. Surveys were avoided on the cloudy, misty, foggy and rainy days. The bird species were identified and recorded using field binocular (8 x 50 Olympus). To facilitate systematic sampling and habitat-wise documentation, a detailed map of the study area was prepared in QGIS 4.0 using high-resolution Google Earth satellite imagery supplemented with field verification (Fig. 1c). A total of 88 field surveys (approximately seven surveys per month) were completed over the 12-month study period. Observations were recorded during the peak periods of bird activity, between 06:00-9:00 h in the morning, and 16:00-18:00 h in the evening, to maximize the probability of visual and acoustic detections. Bird species were identified using standard field guides (Grimmett et al. 2016; Dickinson 2026). A comprehensive checklist of all recorded bird species was prepared, and taxonomic classification followed the latest standardized avian checklists (Maheswaran and Alam 2025; Praveen and Jayapal 2024). Each species was further categorized according to species group, feeding guild, residential status, relative abundance, sighting frequency, global conservation status (IUCN Red List), and global population trend (IUCN 2026).

### Data analysis

Species richness was calculated as the total number of avian species recorded during the study period, and an abundance matrix comprising all observed species was compiled for subsequent analyses. Relative abundance of each species was expressed as the percentage contribution of its total number of individuals to the cumulative abundance of all recorded bird species. Community structure was further examined using a Whittaker rank-abundance plot, in which species were arranged in descending order of their relative abundance to characterize patterns of species dominance and evenness within the avian community (Buckland et al. 2011).

Temporal persistence of individual species was assessed by calculating the sighting frequency, i.e. the percentage of survey days on which a species was recorded relative to the total number of survey days (Helden 2021). Based on their calculated occurrence frequency, species were categorized as abundant (76-100%), common (51-75%), uncommon (26-50%) and rare (0-25%) (MacKinnon and Phillips 1993; Rajashekara and Venkatesha 2017). Alpha diversity was evaluated using diversity, richness, dominance, and evenness indices to characterize community composition across seasons during a year of study (Li N et al, 2019). Sampling completeness and potential species richness were assessed using non-parametric estimators such as, Chao-1, iChao-1 and Abundance-based Coverage Estimator (ACE), each calculated with 95% confidence intervals (Chao A, 1984). All statistical analysis were performed using PAST (Paleontological Statistics, version 5.3) software package to evaluate the taxonomic composition, community structure, and seasonal dynamics of the avian assemblage (Hammer et al. 2001).

## Results

### Habitat composition of the Magadh University campus

We characterized the campus landscape in order to gain spatial information and interpreting any pattern between the distribution and diversity of avifauna recorded during the study period. Habitat classification revealed a heterogeneous mosaic of sparse and dense vegetated areas, open fields, built-up areas and aquatic habitats (Fig. 4). Scattered tree cover constituted the largest habitat category, covering 30.33% of the total campus area, followed by built-up areas (26.20%) and open areas (23.38%). Dense canopy accounted for 19.57% of the campus, whereas water bodies represented a small proportion (0.43%) of the mapped area. Thus, nearly half of the campus was occupied by tree-dominated habitats comprising scattered and dense canopy, while the remaining area consisted of built-up, open habitats and limited aquatic zones. This heterogeneity is likely to provide a range of potential ecological settings for birds with different habitats and resource requirements. Thus, the resulting habitat mosaic formed the basis for assessing the diversity, abundance and seasonal occurrence of the avifauna recorded across the campus.

**Fig. 4.**
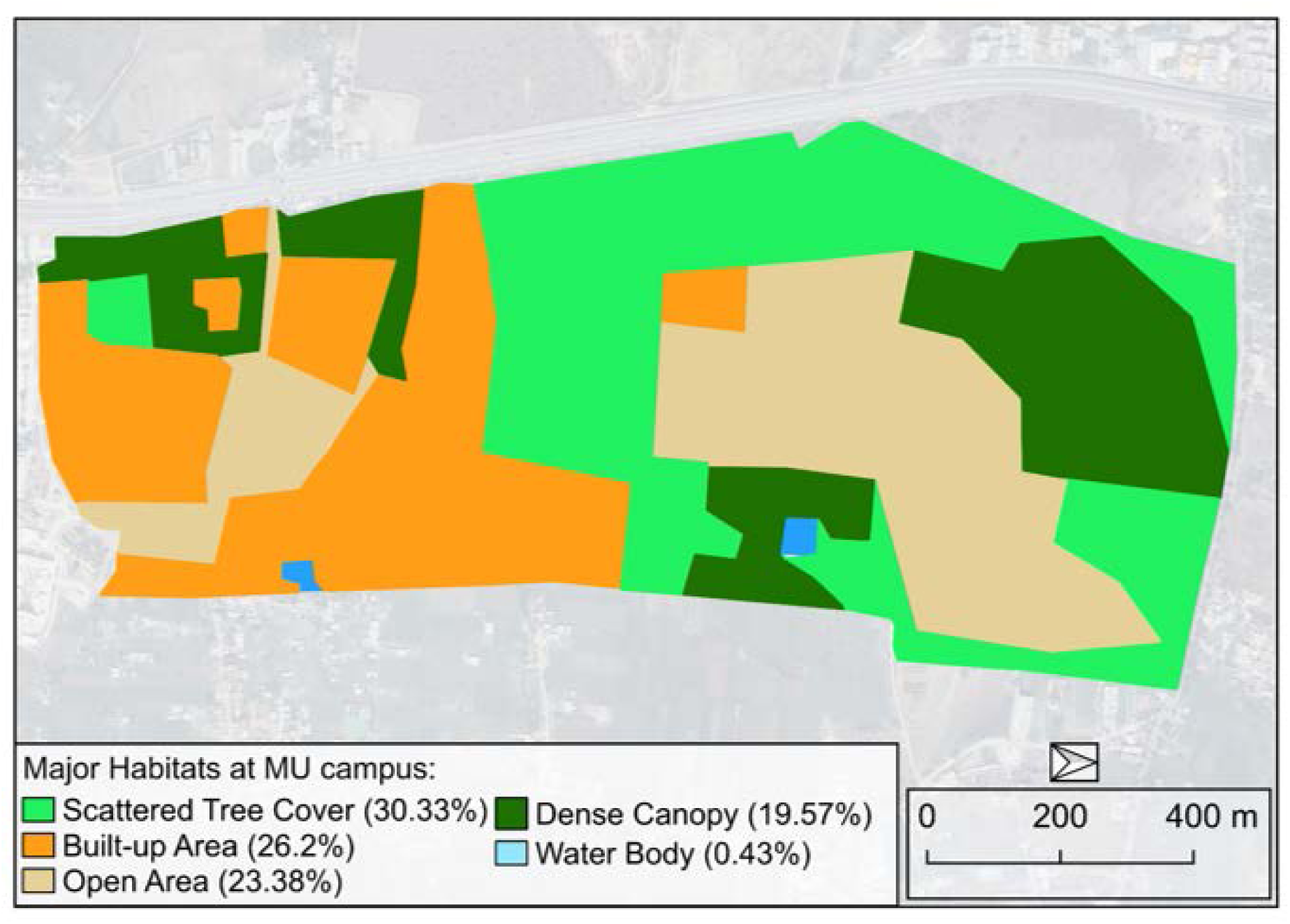
Habitat classification within the MU campus. The QGIS map shows the spatial distribution of five major habitat categories - scattered tree cover, built-up area, open area, dense canopy, and water body, with their proportional contributions to the total campus area.

### Avian diversity of the Magadh University campus

Within this heterogeneous habitat mosaic of the campus, year-round survey documented a diverse avian assemblage representing multiple taxonomic groups. A total of 78 bird species belonging to 41 families and 16 orders were recorded during the 12-month survey period, comprising 37 species groups (Table 1) (Fig. 5). The order Passeriformes was the most species-rich, comprising 37 species *i.e*. 47.44% of the total recorded richness, followed by Coraciiformes (6 species), Columbiformes and Charadriiformes (5 species each), Piciformes and Galliformes (4 species each), and Pelecaniformes and Cuculiformes (3 species each). Strigiformes, Psittaciformes and Apodiformes contributed two species each, whereas Gruiformes, Falconiformes, Ciconiiformes, Bucerotiformes and Accipitriformes were each represented by a single species (Table 2). The hierarchical distribution of avian orders and their constituent families highlighted the pronounced contribution of Passeriformes to the overall taxonomic composition of the campus avifauna (Fig. 5).

**Fig. 5.**
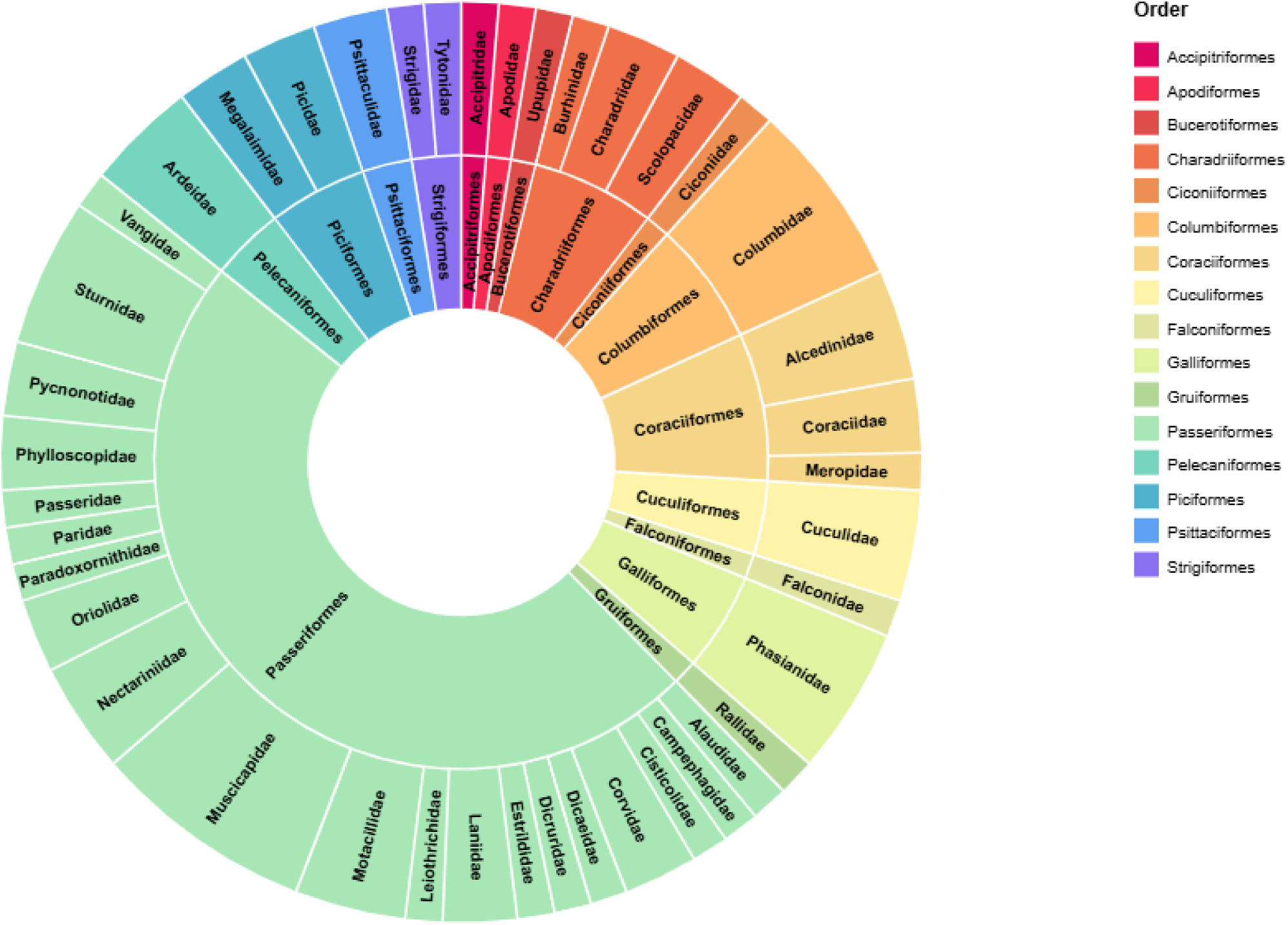
Hierarchical taxonomic composition of the avian community at MU campus. The inner ring represents 16 recorded avian orders, while the outer ring depicts corresponding 41 families within each order. Segment size represents the species richness within each taxonomic group. The figure highlights the richness of Passeriformes within the recorded avian species.

**Table 1.**
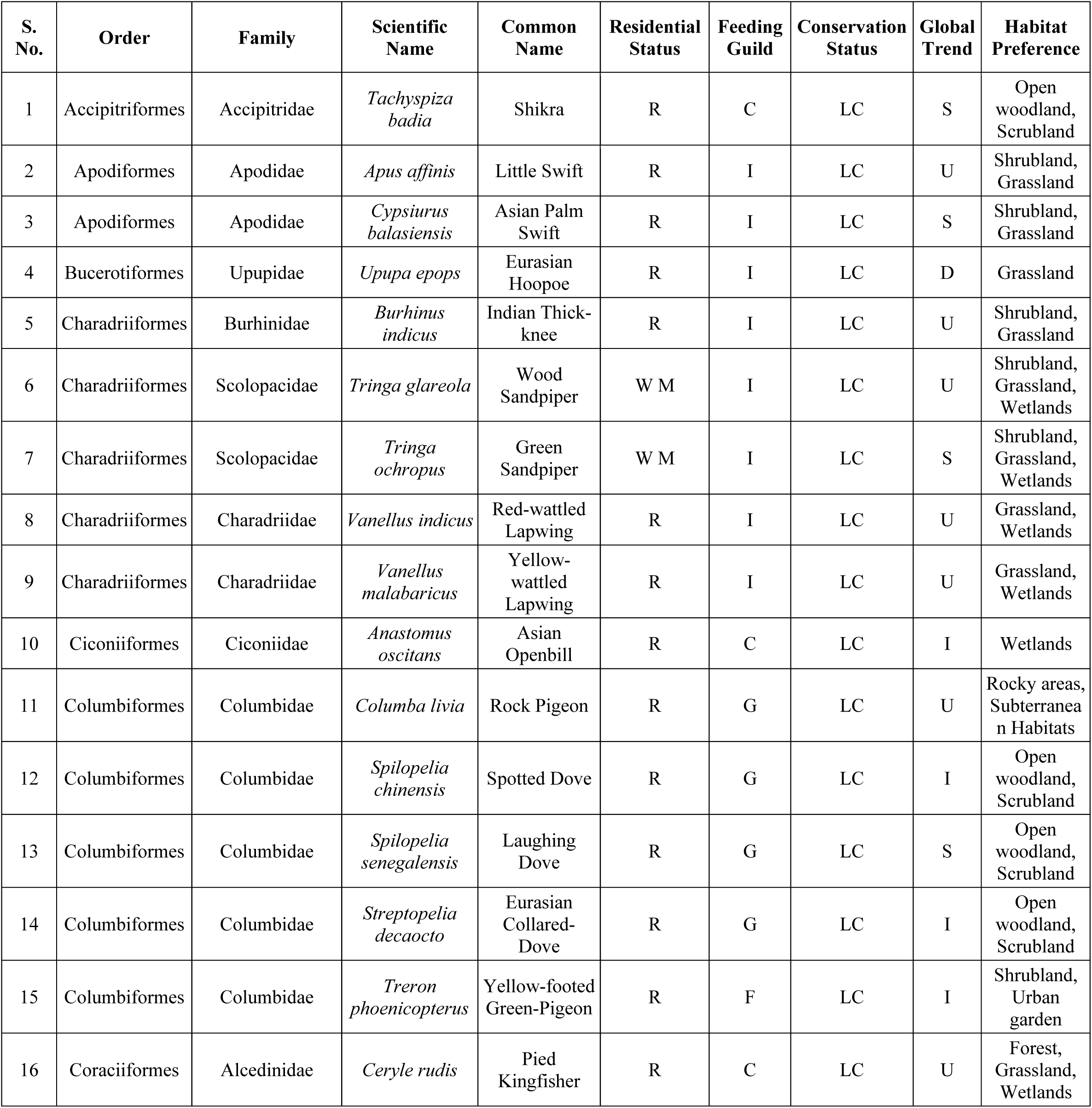
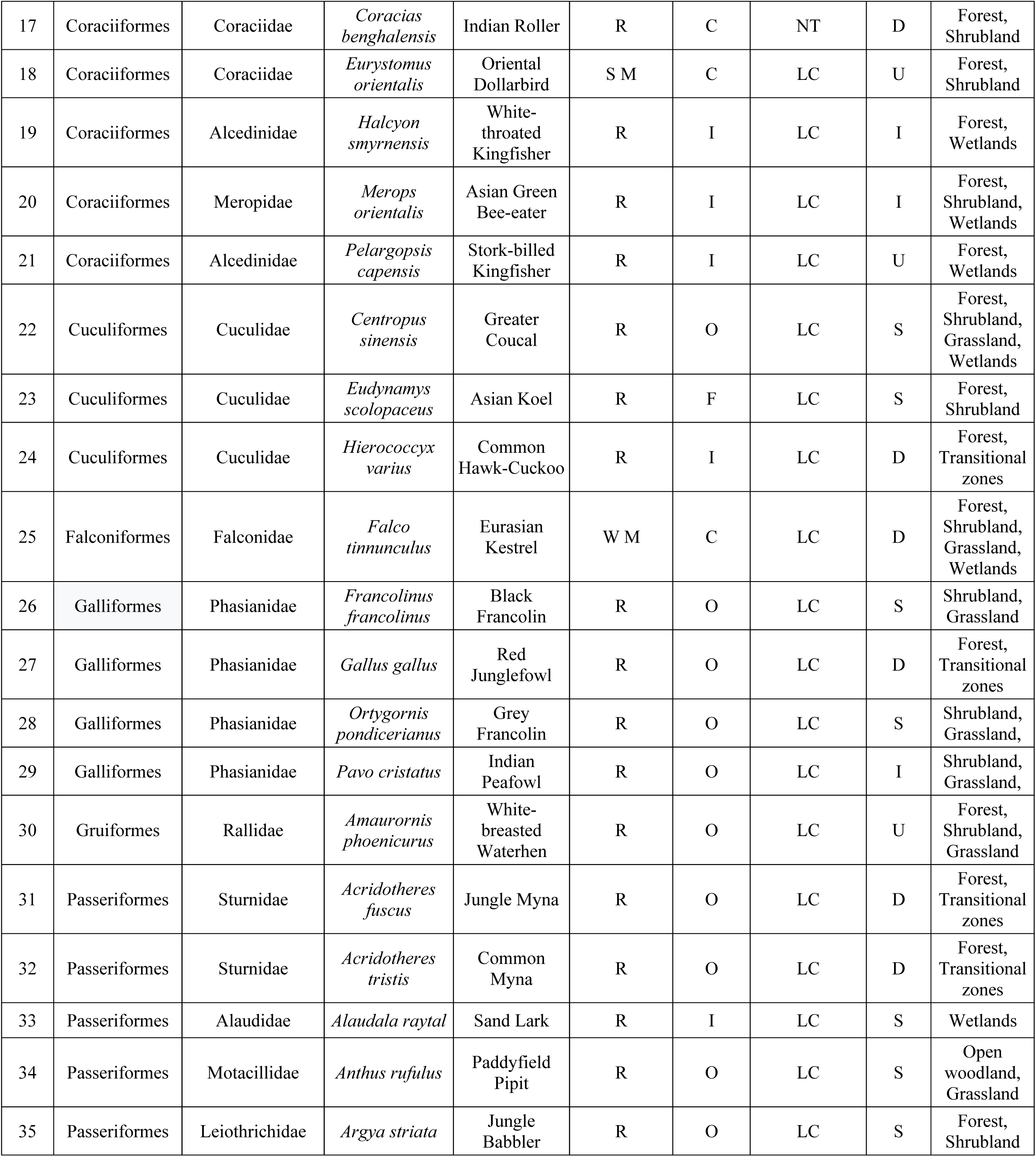
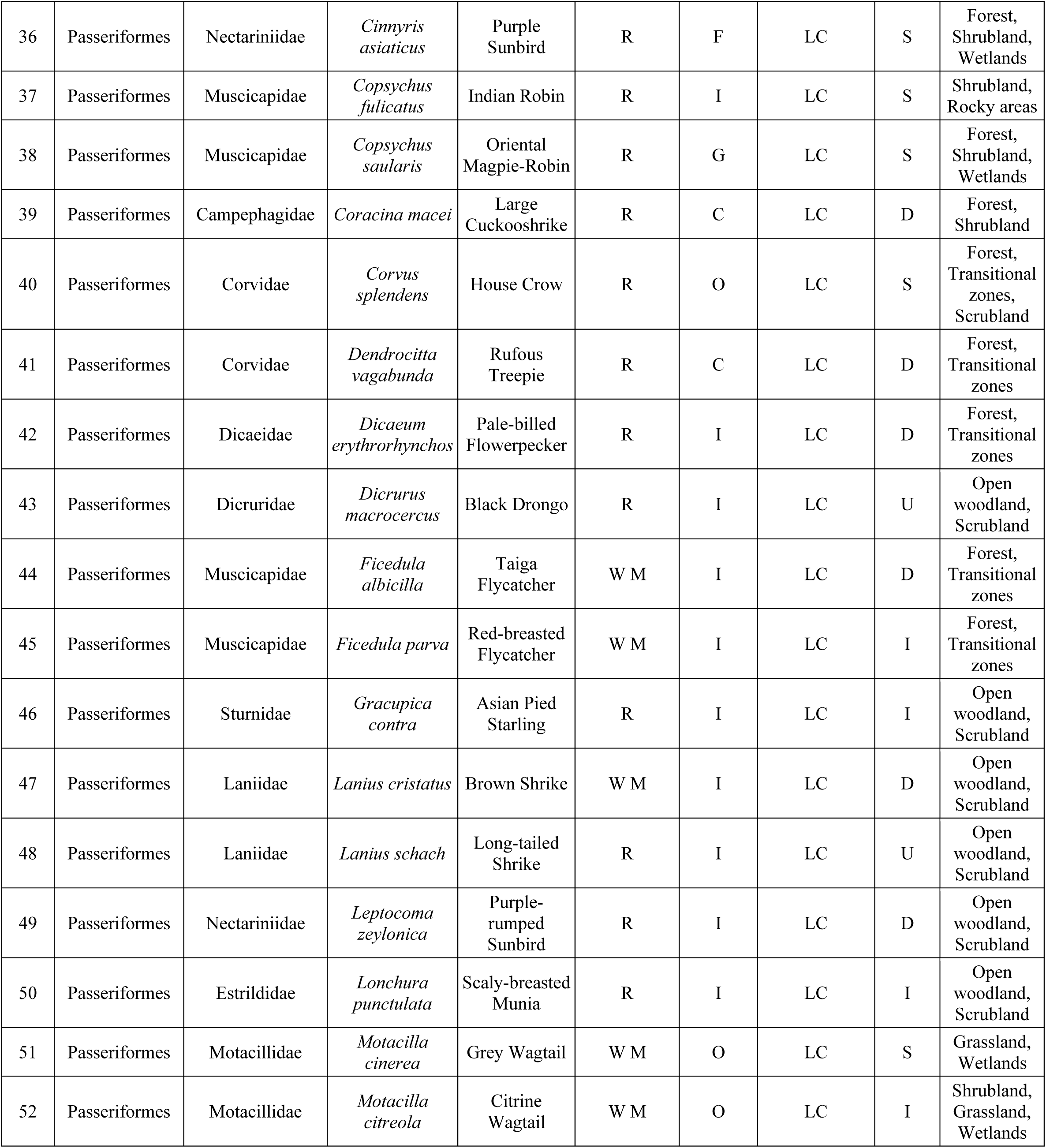
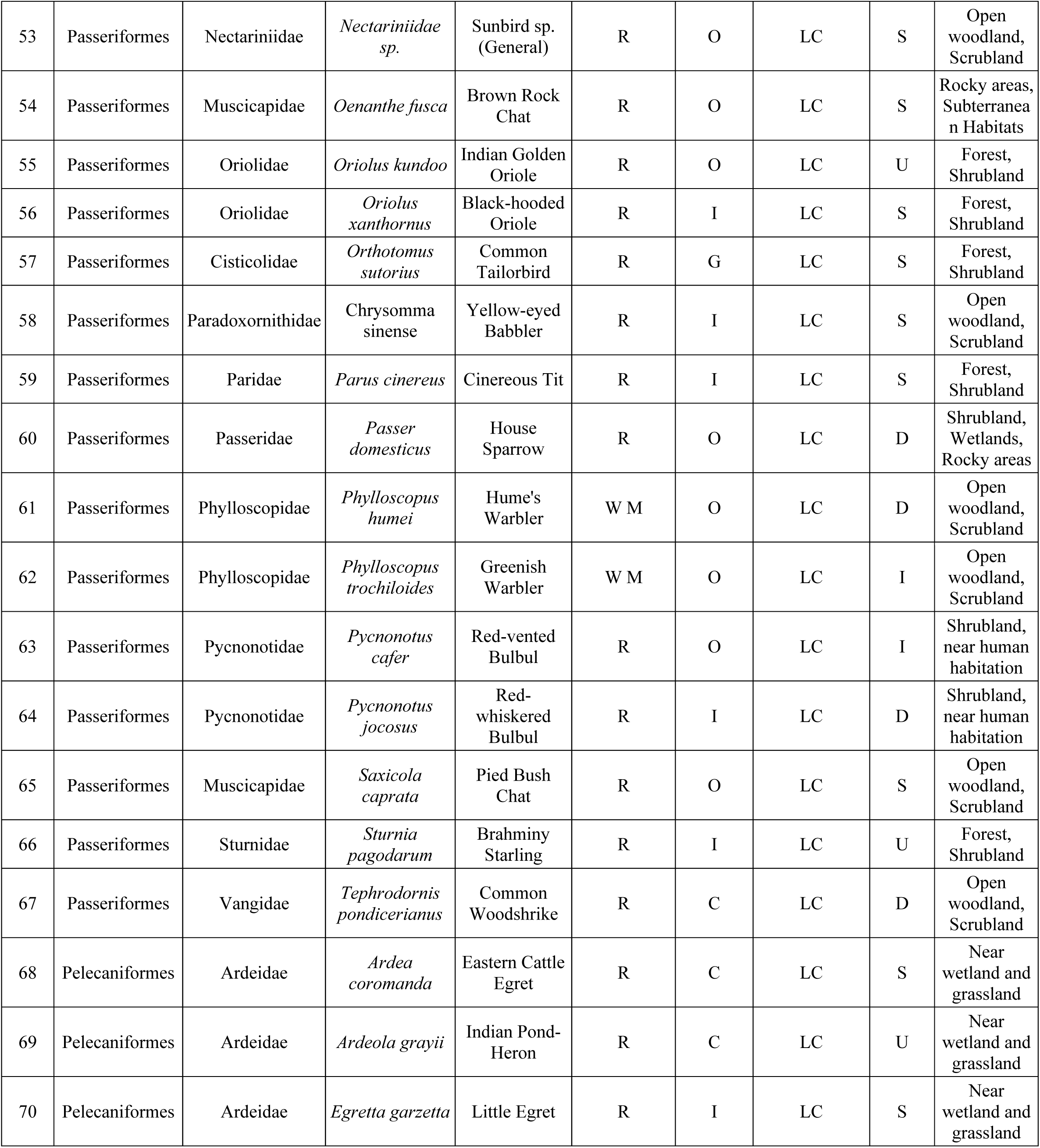
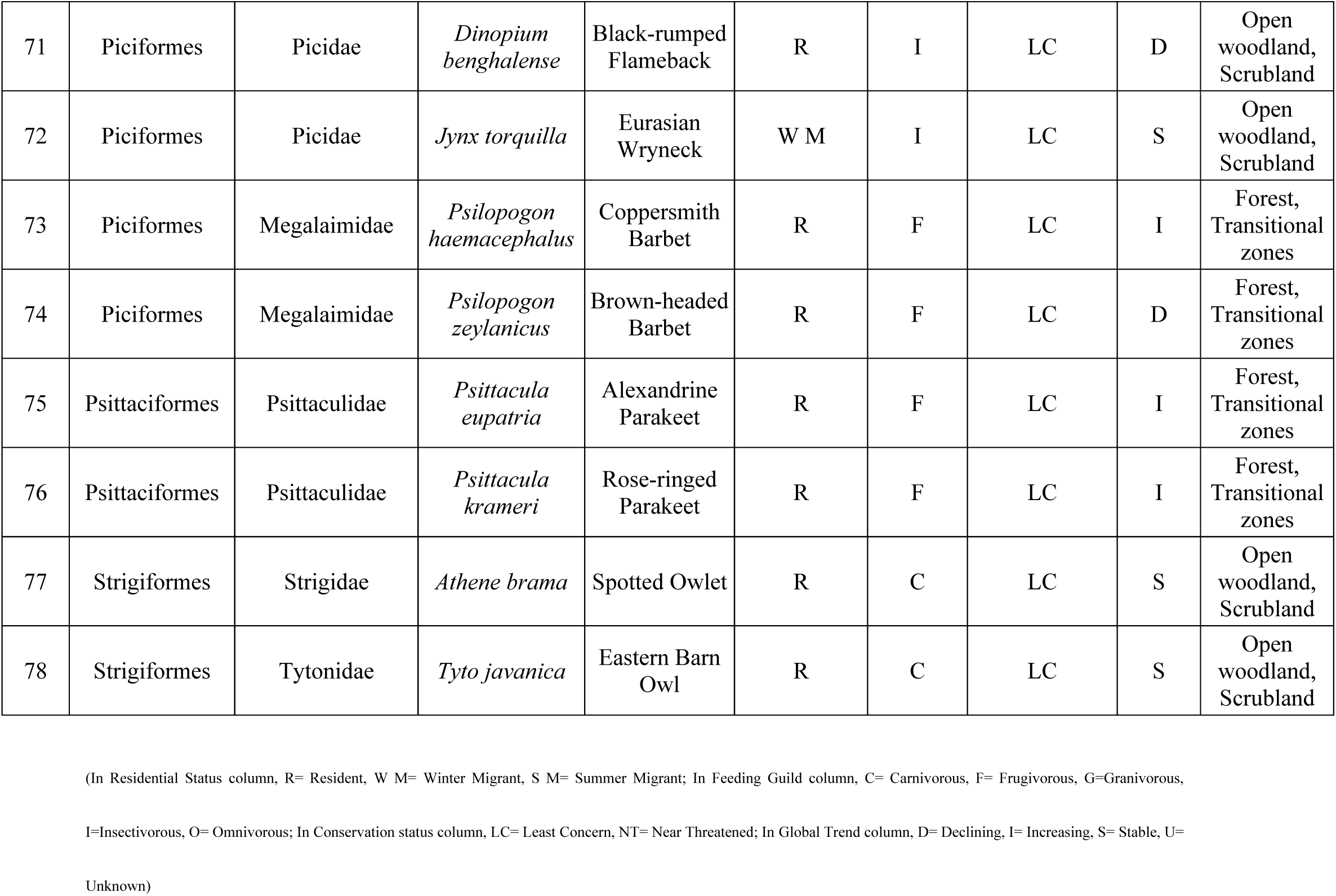
A comprehensive checklist of avian species observed at MU, Bodh Gaya Campus.

**Table 2.**
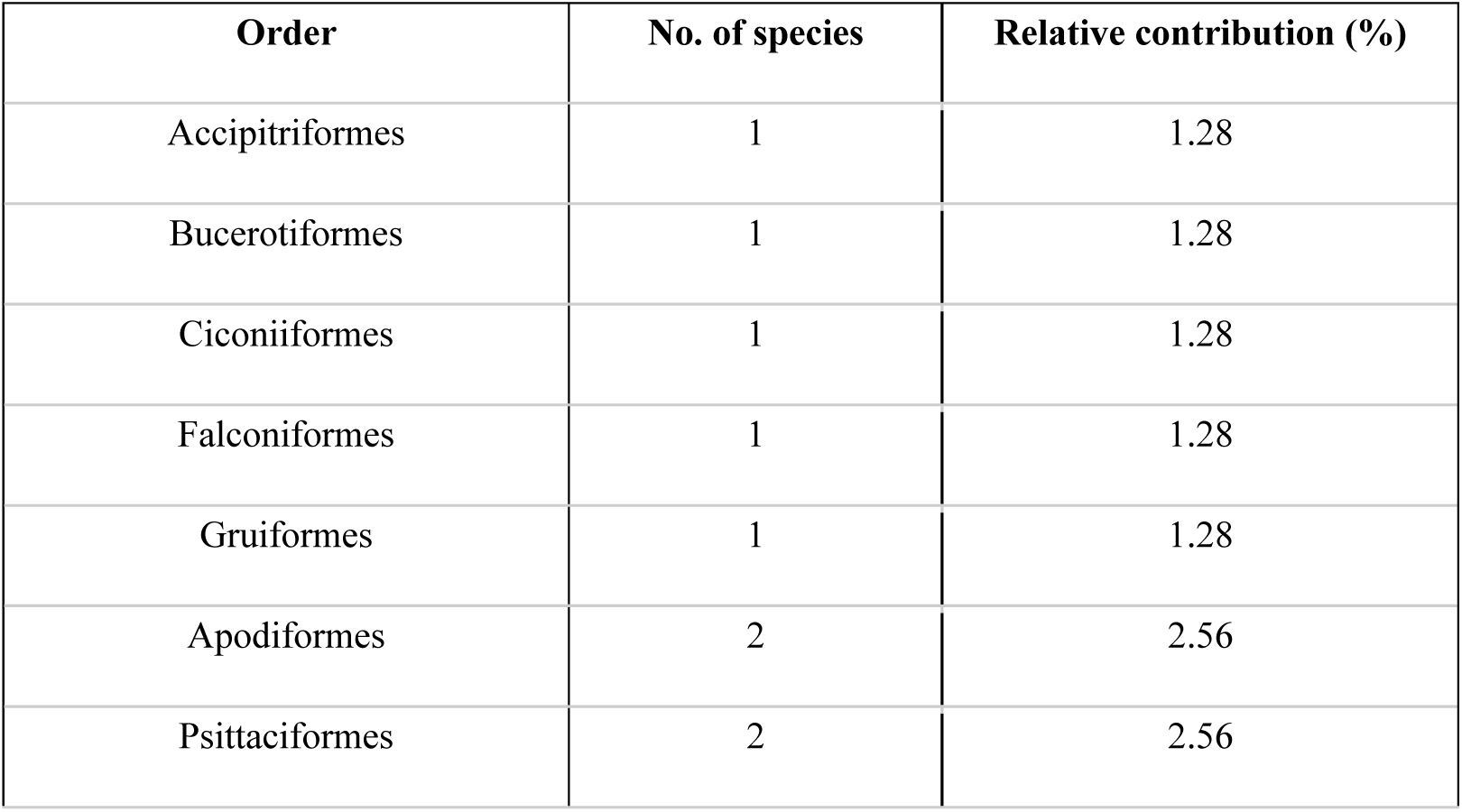
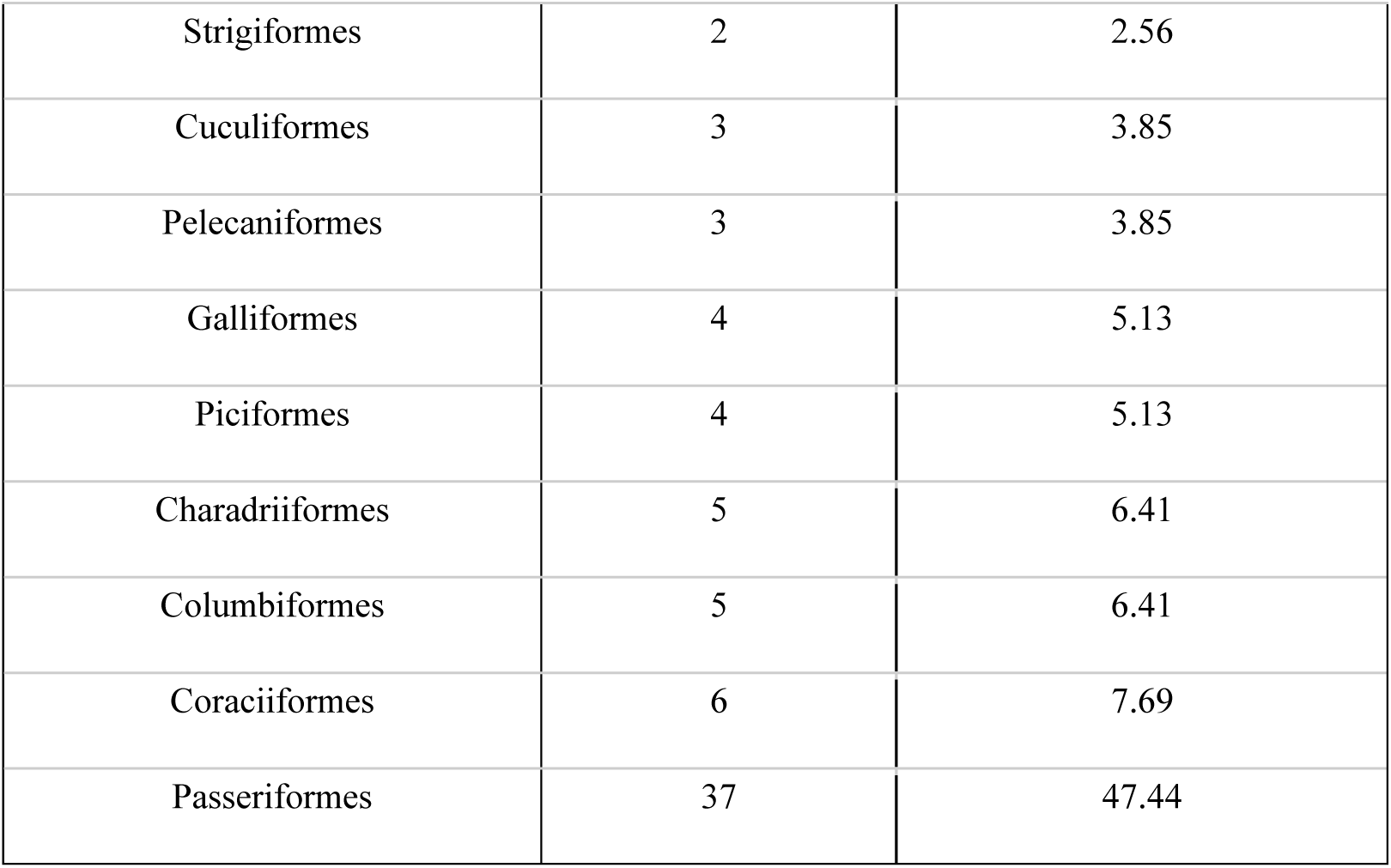
Order-level species richness and relative contribution of avian species at MU campus.

At the family-level, Relative Diversity (RDi) analysis showed that Muscicapidae was the most species-rich family, comprising six species (RDi = 7.69%), followed by Columbidae (5 species, RDi = 6.41%), Phasianidae and Sturnidae (4 species each, RDi = 5.13%) and Alcedinidae, Ardeidae, Cuculidae, Motacillidae and Nectariniidae comprising three species each (RDi = 3.85%). Twelve families – Apodidae, Charadriidae, Coraciidae, Corvidae, Laniidae, Megalaimidae, Oriolidae, Phylloscopidae, Picidae, Psittaculidae, Pycnonotidae and Scolopacidae were represented by two species each (RDi = 2.56%), whereas the remaining 20 families were represented by a single species each (RDi = 1.28%) (Table 3).

**Table 3.**
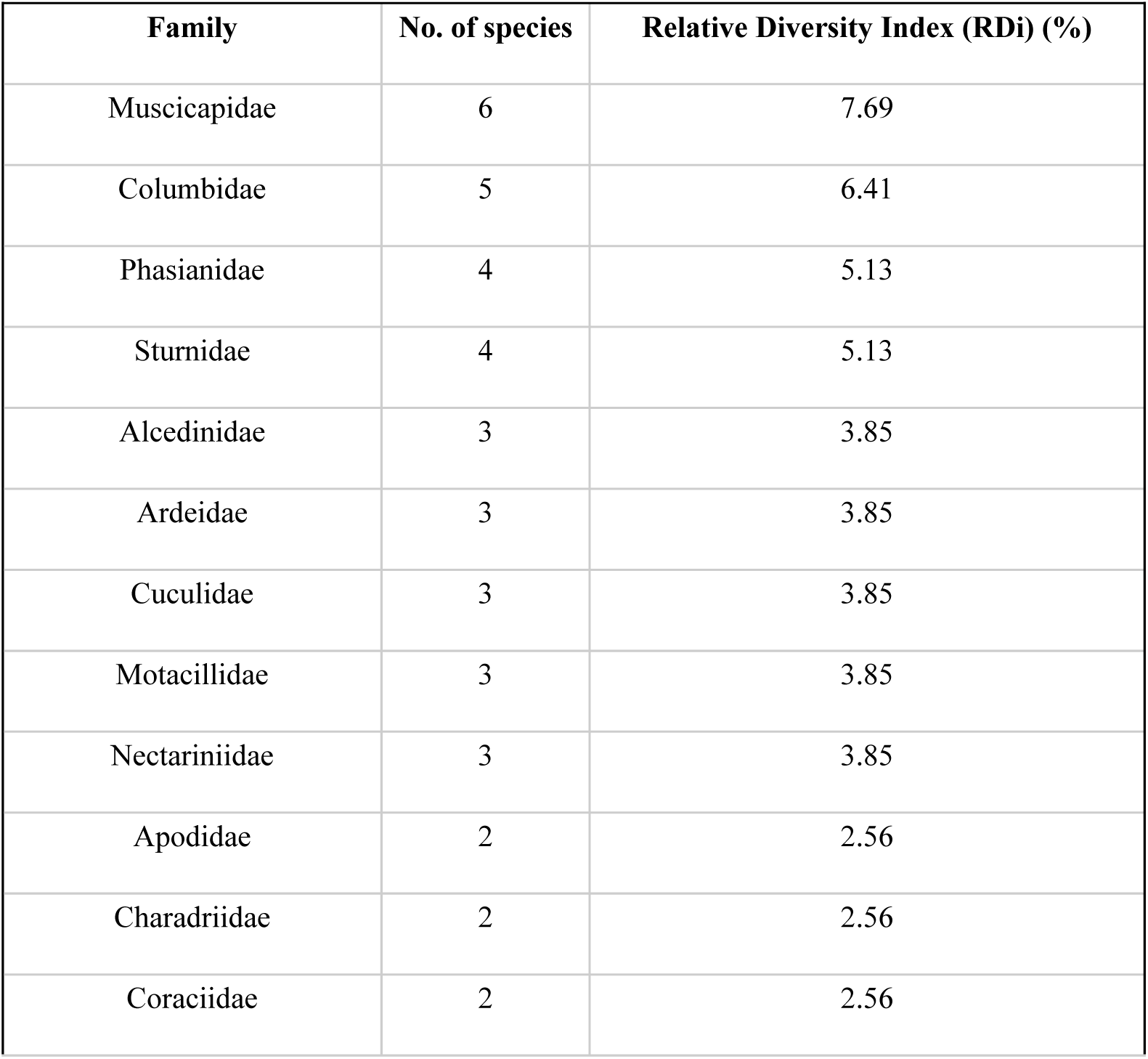
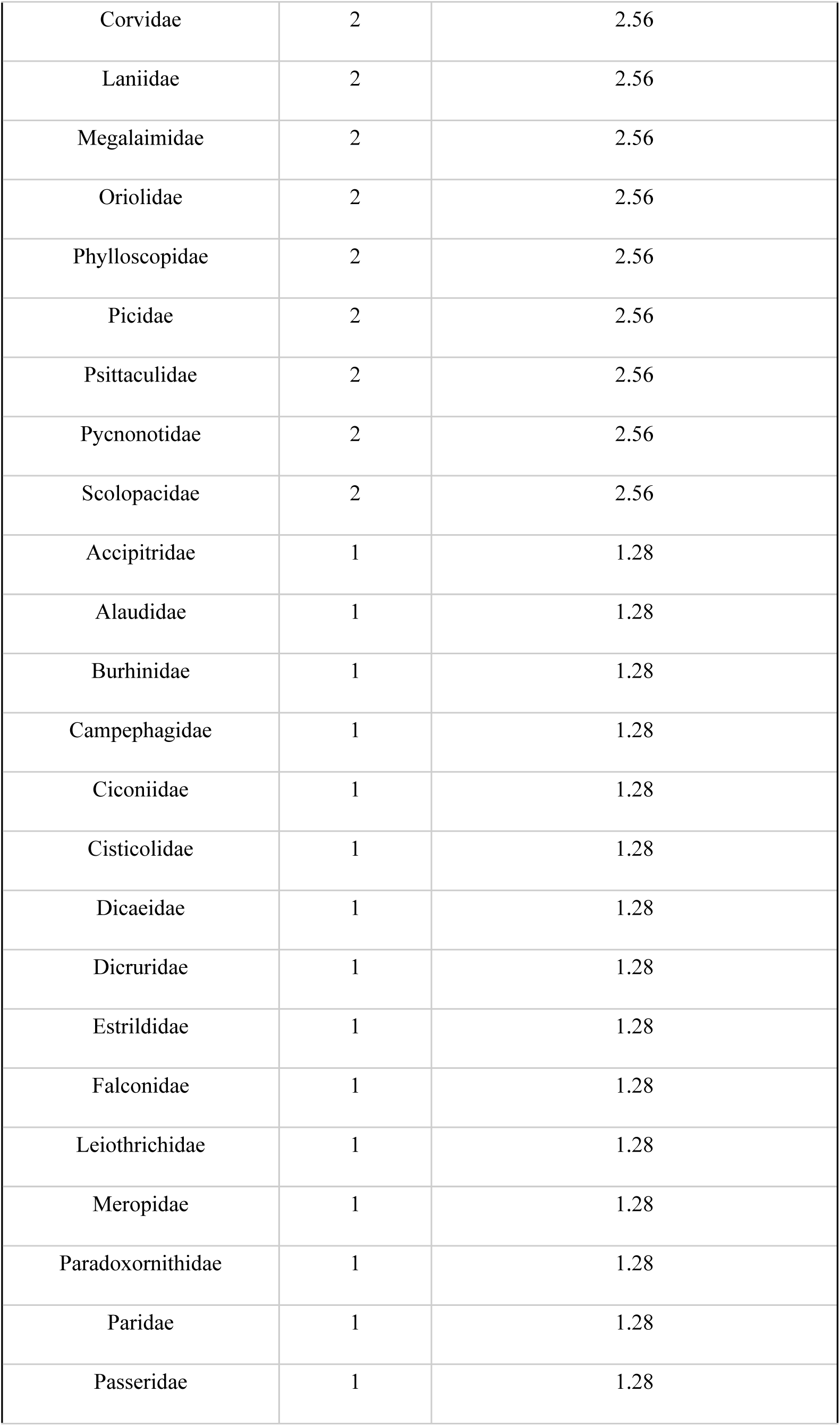
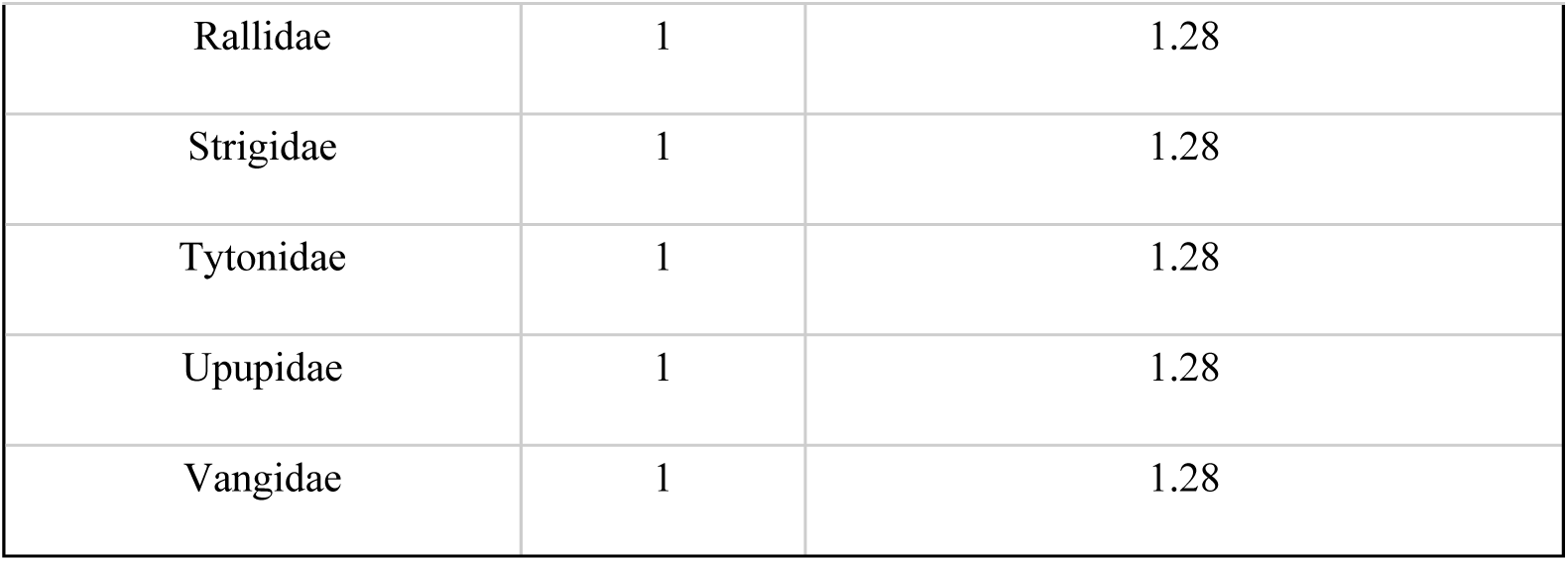
Family-level species richness and Relative Diversity Index (RDi) of avian assemblage at MU campus.

| **Family** | **No. of species** | **Relative Diversity Index (RDi) (%)** |
| --- | --- | --- |
| Muscicapidae | 6 | 7.69 |
| Columbidae | 5 | 6.41 |
| Phasianidae | 4 | 5.13 |
| Sturnidae | 4 | 5.13 |
| Alcedinidae | 3 | 3.85 |
| Ardeidae | 3 | 3.85 |
| Cuculidae | 3 | 3.85 |
| Motacillidae | 3 | 3.85 |
| Nectariniidae | 3 | 3.85 |
| Apodidae | 2 | 2.56 |
| Charadriidae | 2 | 2.56 |
| Coraciidae | 2 | 2.56 |
| Corvidae | 2 | 2.56 |
| Laniidae | 2 | 2.56 |
| Megalaimidae | 2 | 2.56 |
| Oriolidae | 2 | 2.56 |
| Phylloscopidae | 2 | 2.56 |
| Picidae | 2 | 2.56 |
| Psittaculidae | 2 | 2.56 |
| Pycnonotidae | 2 | 2.56 |
| Scolopacidae | 2 | 2.56 |
| Accipitridae | 1 | 1.28 |
| Alaudidae | 1 | 1.28 |
| Burhinidae | 1 | 1.28 |
| Campephagidae | 1 | 1.28 |
| Ciconiidae | 1 | 1.28 |
| Cisticolidae | 1 | 1.28 |
| Dicaeidae | 1 | 1.28 |
| Dicruridae | 1 | 1.28 |
| Estrildidae | 1 | 1.28 |
| Falconidae | 1 | 1.28 |
| Leiothrichidae | 1 | 1.28 |
| Meropidae | 1 | 1.28 |
| Paradoxornithidae | 1 | 1.28 |
| Paridae | 1 | 1.28 |
| Passeridae | 1 | 1.28 |
| Rallidae | 1 | 1.28 |
| Strigidae | 1 | 1.28 |
| Tytonidae | 1 | 1.28 |
| Upupidae | 1 | 1.28 |
| Vangidae | 1 | 1.28 |

Community-level diversity analysis indicated relatively high avian diversity and low overall dominance. The Shannon-Wiener diversity index was H’ = 3.278, while Simpson’s dominance value was low (D = 0.062), corresponding to a high Simpson’s diversity value (1-D = 0.938). Together, these indices indicate that the recorded individuals were distributed across a relatively diverse assemblage without pronounced dominance by a small number of species. Pielou’s evenness J = 0.751 indicates that the abundance is reasonably even. The Margalef richness index (8.472) and Menhinick richness index (0.829) further characterized the species-rich nature of avian assemblage within the MU campus. The Berger-Parker dominance index was 0.118, indicating that the most abundant species accounted for 11.8% of total recorded abundance and that the assemblage lacked strong dominance by a single species (Table 4).

**Table 4.**
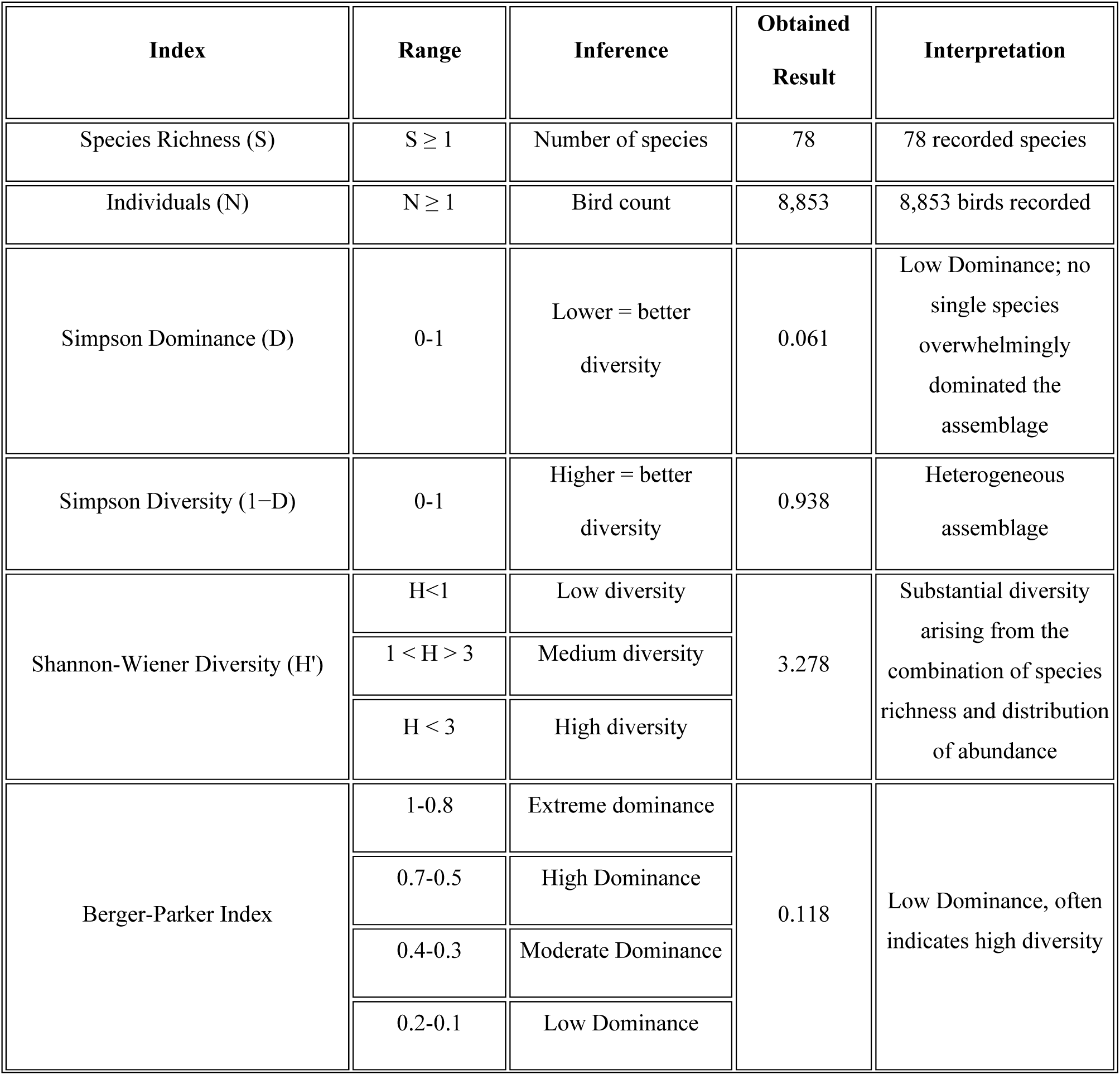
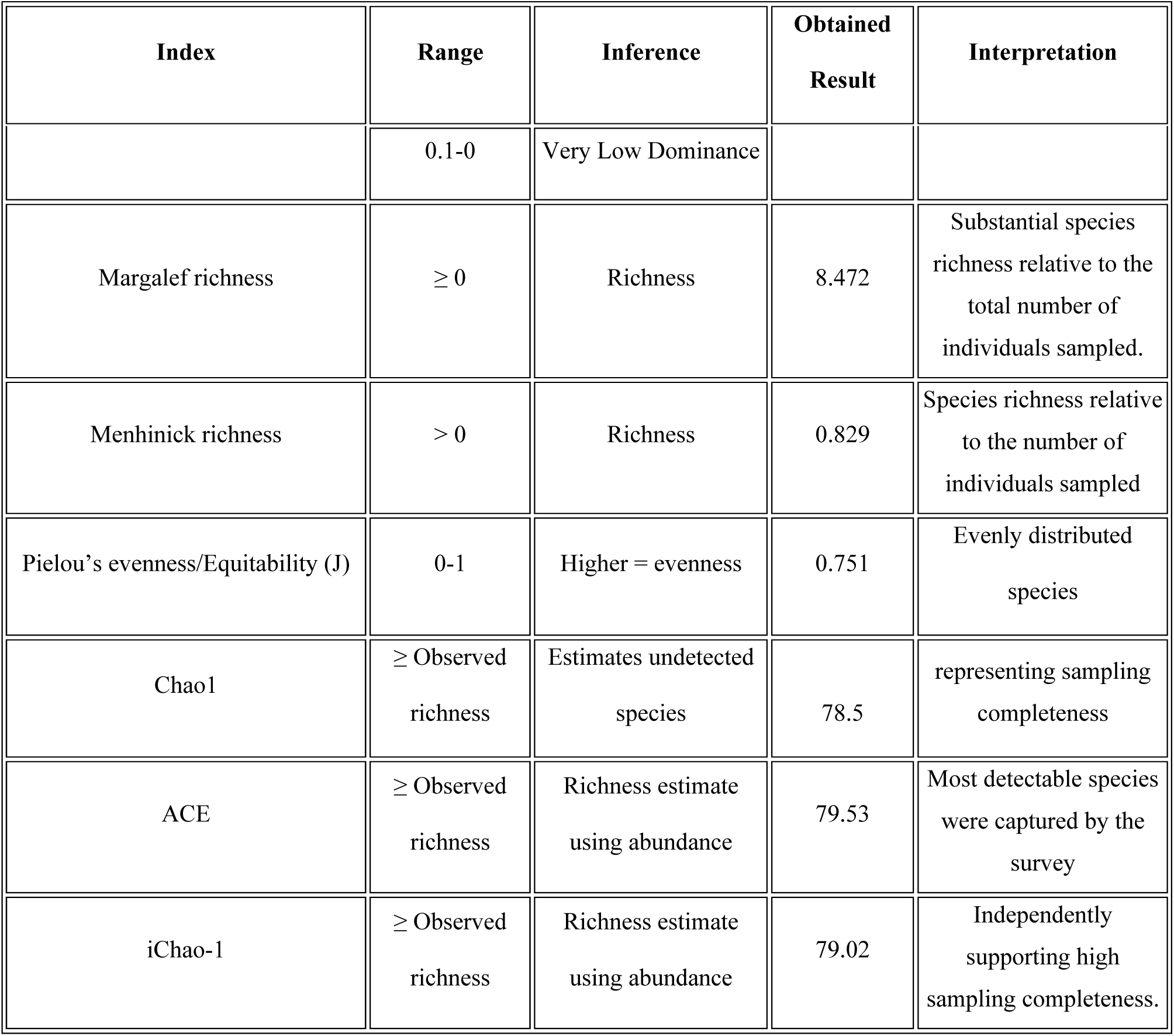
Community-level diversity indices and richness estimates of avian assemblages at MU campus.

Sampling completeness was evaluated using non-parametric richness estimators. The estimated species richness obtained from Chao1 (78.5 species), iChao-1 (79.02) and the Abundance-based Coverage Estimator (ACE; 79.53 species) closely matched the observed richness of 78 species. Furthermore, the species accumulation curve approached an asymptote, indicating a declining rate of new species detection with increasing sampling effort in the study area. (Fig. 6).

**Fig. 6.**
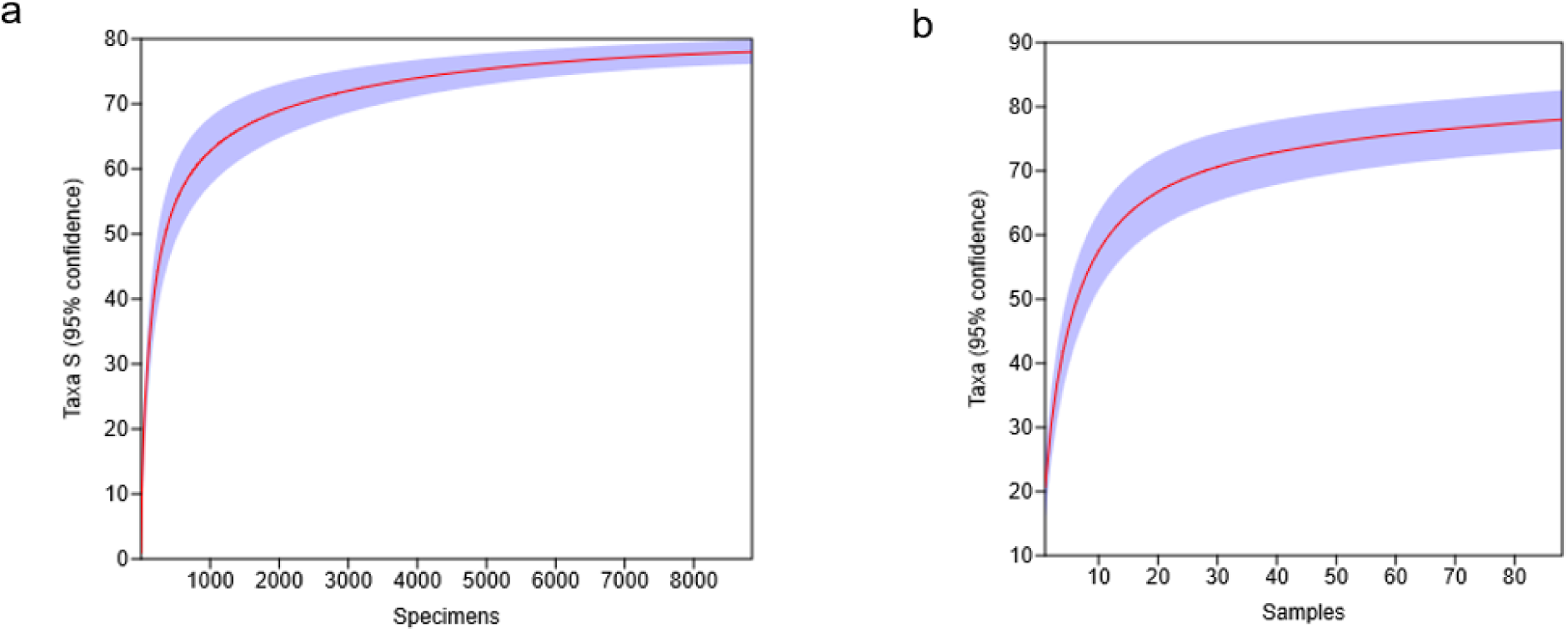
Assessment of sampling completeness of the avian assemblage at MU campus. (a) Individual rarefaction curve based on annual abundance of birds at MU campus. (b) Sample-based rarefaction curve (Mao’s Tau) based on the 88 survey occasions. The blue-shaded area shows the 95% confidence interval.

### Species abundance and community structure

The recorded avian species differed in their contribution to the overall number of individuals, indicating an uneven abundance within campus assemblage. *Argya striata* (Jungle Babbler) was the most abundant species, accounting for 11.79% of the total individuals recorded, followed closely by *Acridotheres tristis* (Common Myna; RA = 9.94%)*, Spilopelia chinensis* (Spotted Dove; RA = 9.94%), *Pycnonotus cafer* (Red-vented Bulbul; RA = 9.04%), and *Cinnyris asiaticus* (Purple Sunbird; RA = 8.74%). In contrast, several species contributed less than 1% to the total recorded abundance (Fig. 7).

**Fig. 7.**
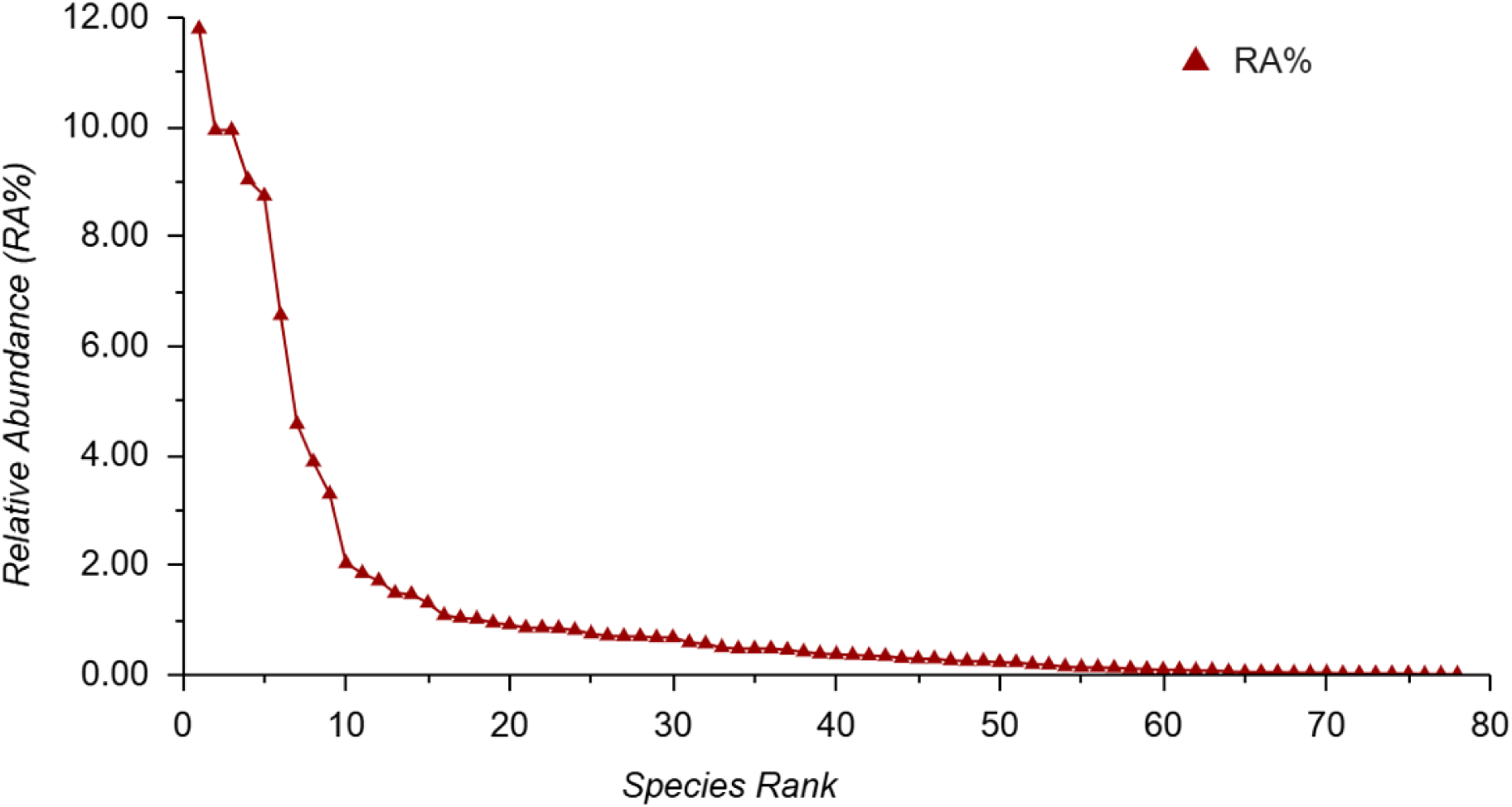
Rank-abundance distribution of the avian species recorded at MU campus. Species are arranged in descending order of relative abundance, showing the pattern of individual abundance among recorded species.

The rank-abundance curve showed a sharp decline in relative abundance among the highest-ranked species, followed by a prolonged tail comprising of numerous species with comparatively low abundance (Fig. 7). This pattern indicates that abundance was concentrated among a relatively small subset of species, while a larger proportion of the recorded species occurred at low abundance. Thus, the campus supported considerable species richness, but the individuals were not distributed evenly across the recorded species.

### Feeding guild and habitat association of recorded avian assemblage

The recorded avian assemblage comprised species with diverse dietary habits, reflecting broad variation in the resource use across the campus habitats. Based on their primary feeding preferences, the 78 recorded species were classified into five major feeding guilds - insectivorous, omnivorous, carnivorous, frugivorous and granivorous (Fig. 8). Insectivorous species formed the largest guild, comprising 31 species (39.74%), followed by omnivorous with 21 species (26.92%), carnivorous with 13 species (16.67%), frugivorous with 7 species (8.97%), and granivorous with 6 species (7.69%).

**Fig. 8.**
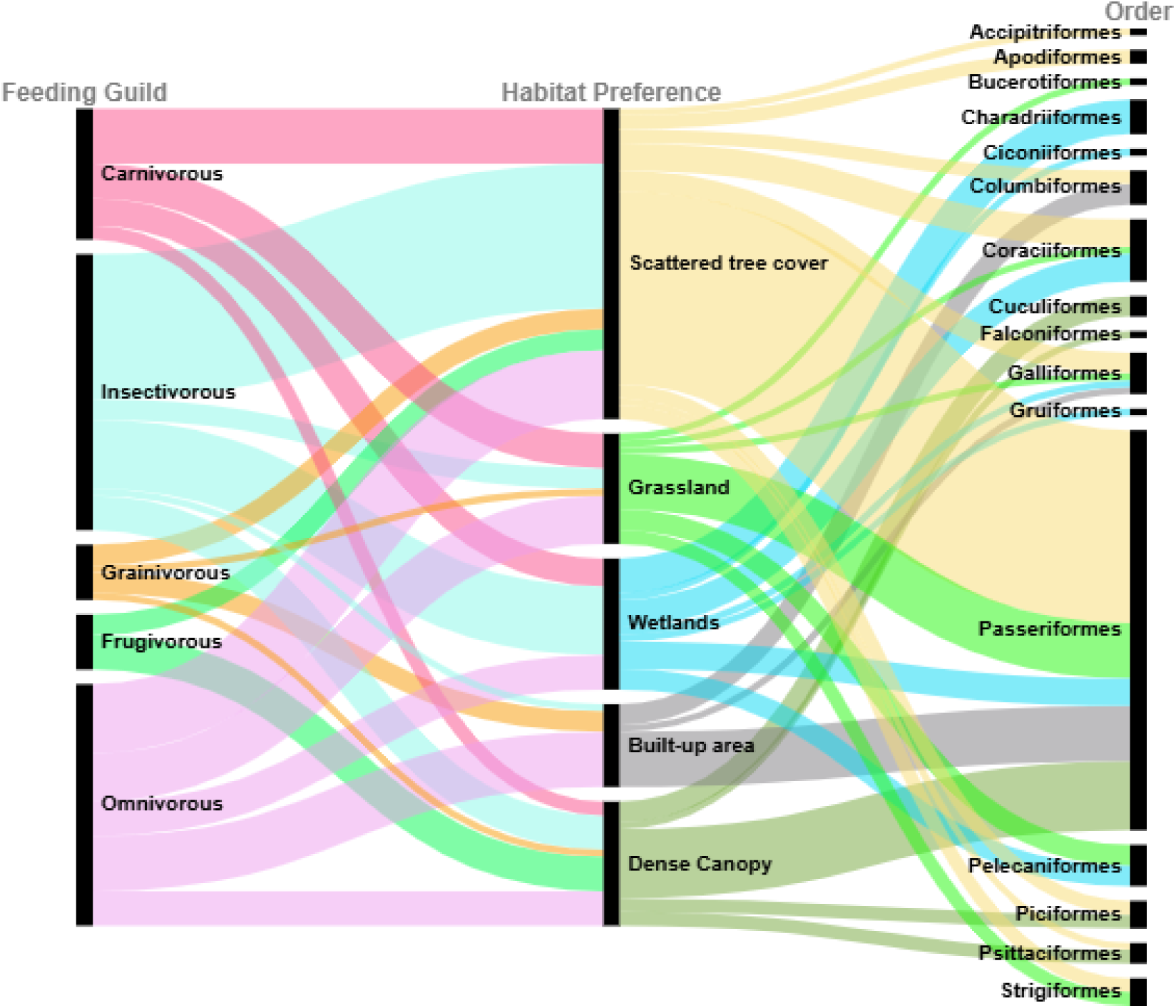
Association between the feeding guilds, habitat preference of the avian assemblage at MU campus. Plot showing the distribution of recorded avifaunal species among five major feeding guilds with their habitat association and their corresponding taxonomic orders. The left side represents various feeding guilds, central part represents their habitat preferences, while the right side represents the recorded avian orders at MU campus. The width of each connecting flow represents number of species assigned to the respective order-feeding-habitat preference combination. Scattered tree cover appeared to be the most preferred habitat.

The Alluvial diagram illustrates the association between the recorded avian orders, their feeding guilds and habitat preference at the study site. Avian assemblage at MU campus comprised five major feeding guilds-carnivorous, insectivorous, omnivorous, frugivorous and granivorous distributed across five major habitat types (Fig. 8). Insectivorous (31) species formed the largest feeding guild followed by omnivorous (21), carnivorous (13), frugivorous (7) and granivorous (6). Insectivorous and omnivorous species were found exploiting all 5 habitat types under study while the remaining showed relatively fewer feeding-habitat association. Scattered tree cover showed serving the species from majority of the avian orders, belonging to diverse feeding guilds. Passeriformes was the most prominent taxonomic order representing multiple habitat preferences and feeding guilds. Other orders showed more restricted or specialized associations with particular feeding guilds.

### Residential status of the recorded avifauna

Based on their established occurrence patterns during the study period, the recorded avian species exhibited three residential categories - resident, winter migrant and summer migrant. Resident species constituted the largest component of the avian community, comprising 66 species (84.62%) of the total recorded richness, whereas 11 species (14.10%) were classified as winter migrants, and one species (1.28%), *Eurystomus orientalis*, was classified as a summer migrant (Fig. 9). The winter migrant species included *Tringa glareola, Tringa ochropus, Falco tinnunculus, Ficedula albicilla, Ficedula parva, Lanius cristatus, Motacilla cinerea, Motacilla citreola, Phylloscopus humei, Phylloscopus trochiloides* and *Jynx torquilla*. Thus, the recorded avian assemblage in the campus predominantly composed of resident species, with a smaller representation of winter and summer migrants.

**Fig. 9.**
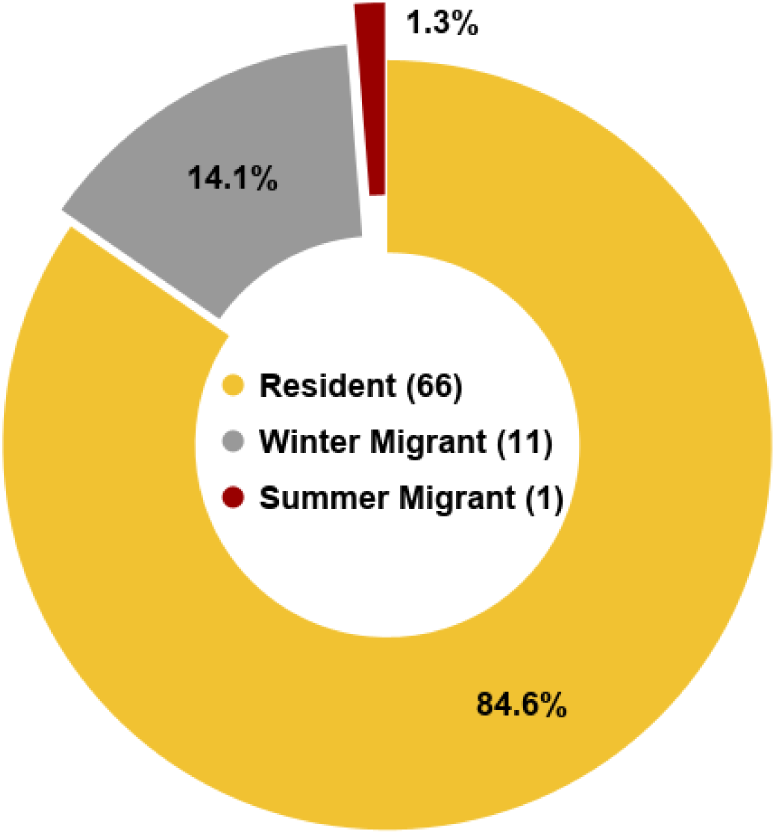
Residential status of avian community at MU campus. The chart shows proportion of resident (84.6%) species, winter migrant (14.1%) species, and one summer migrant (1.3%) found within the campus area during the survey period.

### Seasonal occurrence and annual persistence of the avian community

Seasonal occurrence and annual persistence were assessed to characterize variation in the sighting frequency and temporal distribution of bird species across the study period. Based on sighting frequency, the species were categorized as abundant (100-76%), common (75-51%), uncommon (50-26%) and rare (25-0%). Out of 78 recorded species, 9 species were abundant, 3 species were common, 15 species were uncommon and 51 were of rare occurrence during the year (Fig. 10a). Thus, the campus avian assemblage comprised a relatively small core of frequently detected species, along with a larger group of seasonally or sporadically occurring taxa. Abundant species with occurrence frequency above 75% indicates that the campus provides a baseline of necessary resources *e.g.* nesting sites, year-round food availability, stable habitat *etc*. that can sustain avian life across different seasons.

**Fig. 10.**
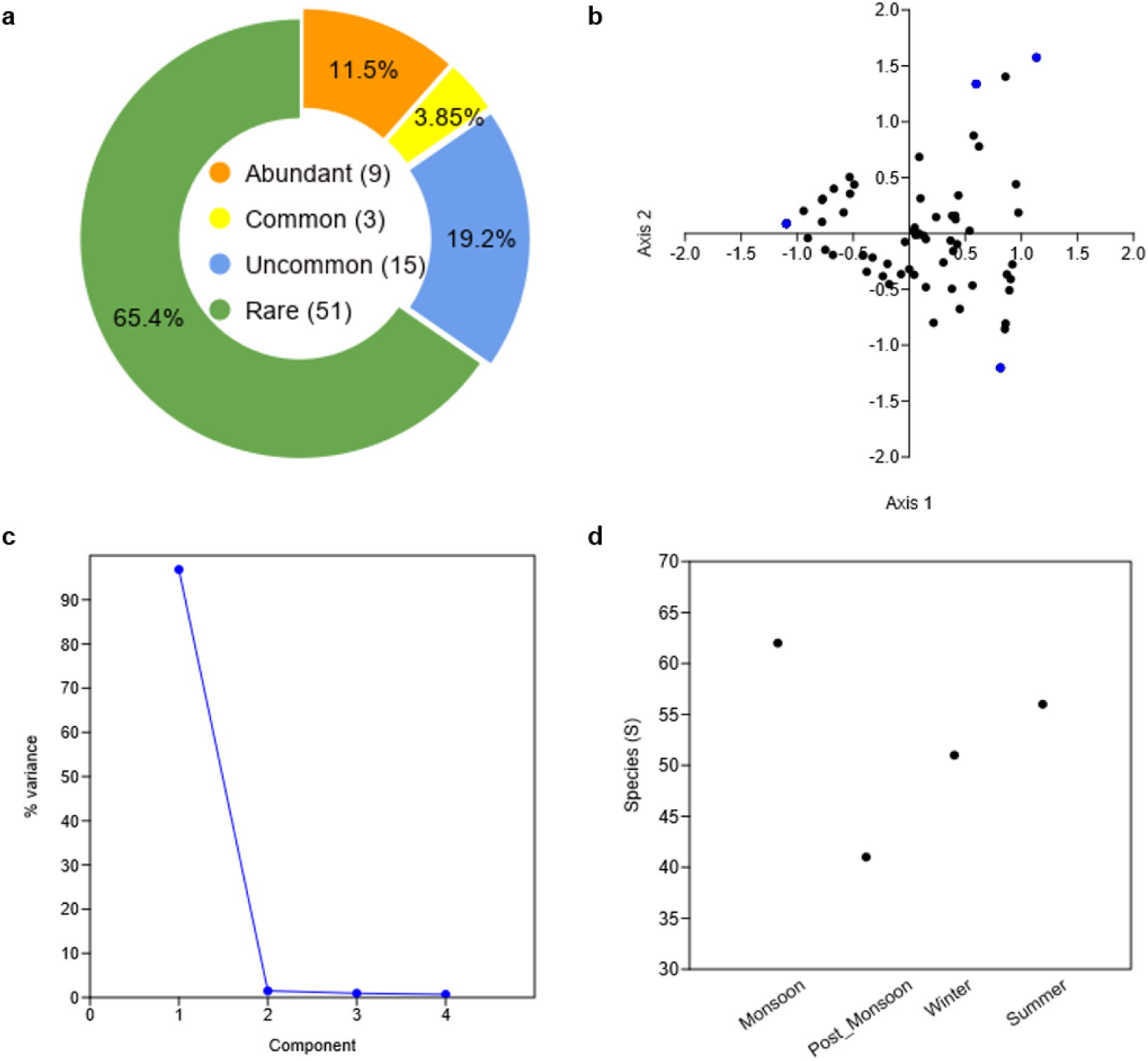
Annual occurrence and community structure of avifaunal assemblage at MU campus. (a) Proportion of species in the four sighting frequency categories - Abundant, Common, Uncommon and Rare based on the sighting frequency throughout the study period. (b) Correspondence Analysis (CA) ordination showing association between recorded bird species and four seasons. Avian species and four seasons are shown as black and blue dots, respectively. (c) Principal Component Analysis (PCA) scree plot showing the percentage of variance based on correlation matrix explained by successive principal components. PC1 and PC2 accounted for 96.836% and 1.503% of the total variance, respectively, and, PC3 and PC4 account for 0.954% and 0.707% of the total variance, respectively. (d) Seasonal species richness for the avian assemblage during the four seasons. Monsoon (62 species) showed the highest richness among the 4 studies seasons and post-monsoon had the lowest richness (41 species)

Correspondence Analysis (CA) demonstrated the relationship between species occurrence and four seasons (Fig. 10b). Most species were positioned close to the centre of the ordination plot, whereas smaller number of species occurred closer to individual seasonal points indicating varying degrees of association with particular seasons. The four seasonal points were positioned relatively close, while species showed varying degree of proximity to the respective seasonal positions. The ordination therefore showed both broadly distributed species and species with stronger seasonal associations.

Principal Component Analysis (PCA) using correlation matrix showed percent variation in bird abundance. PC1 and PC2 accounted for 96.836% and 1.503% of the total variance, respectively, together representing 98.339% of the total variance (Fig. 10c). The scree plot showed a sharp decline from PC1 to PC2, followed by a shallow decline in subsequent components PC3 (0.954) and PC4 (0.706), indicating that most of the variation was represented by the first two components.

Seasonal Simpson’s dominance (D) values showed relatively little variation among the four seasons, ranging from 0.055 in the monsoon season to 0.071 during post-monsoon (Fig. 10d). Post-monsoon season recorded the highest seasonal Simpson dominance value, whereas monsoon recorded the lowest. The winter and summer seasons showed intermediate values 0.069 and 0.068, respectively. Overall, the seasonal dominance estimates show less variation across the four surveyed seasons.

Taken together, these analyses documented variation in the frequency, seasonal association, and abundance of individual species, while the ordination and dominance showed limited separation among the four seasonal assemblages (Verma and Murmu 2015).

### Conservation status of the recorded avifaunal species

The recorded avifauna was further evaluated in terms of global conservation status and population trends to characterize the conservation significance of the species assemblage. The conservation assessment based on the IUCN Red List revealed that 77 of the 78 recorded species (98.7%) were classified as Least Concern (LC), whereas *Coracias benghalensis* (Indian Roller) was categorized as Near Threatened (NT), representing 1.3% of the recorded species (Fig. 11). Despite the predominance of species in the LC category, 19 species were associated with declining global population trends. These findings indicate that, although most species currently have relatively low global conservation concern, the campus supports taxa for which continued habitat protection and long-term population monitoring may be important.

**Fig. 11.**
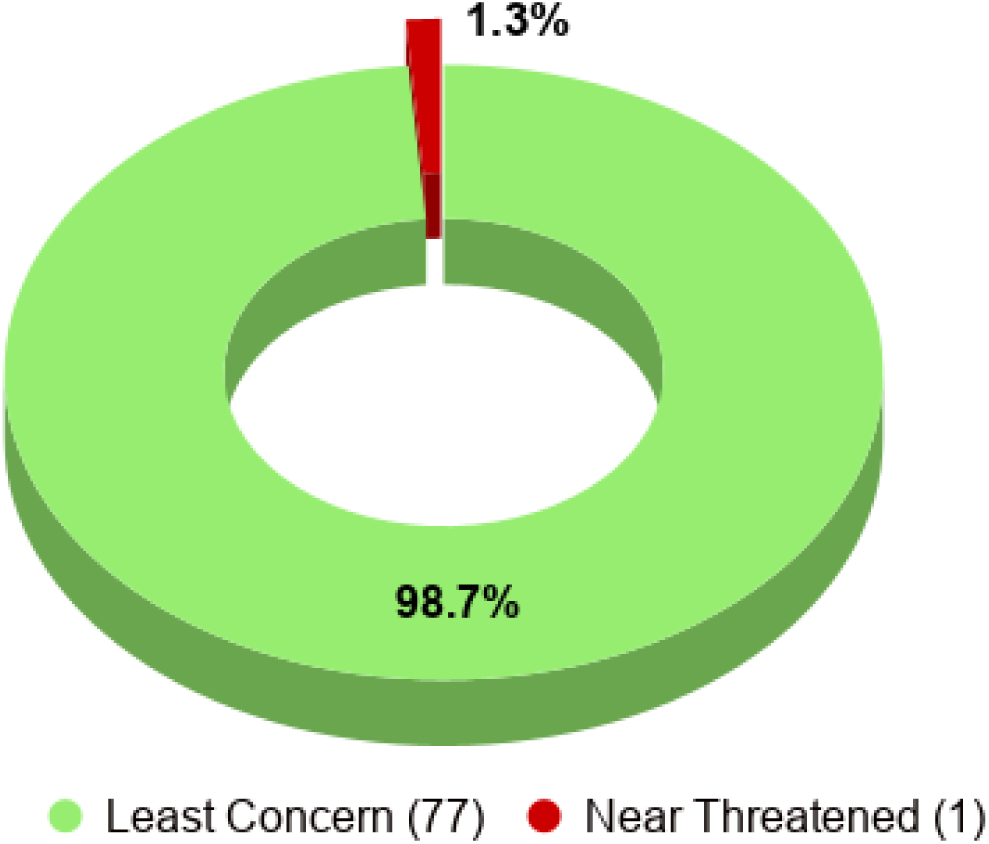
IUCN Red List conservation status of the avian species at MU. The chart depicts the proportion of species according to their IUCN conservation status, with 77 species (98.7%) classified as Least Concern (LC) and one species (1.3%), *Coracias benghalensis*, classified as Near threatened (NT).

## Discussion

The present study documented 78 avian species belonging to 41 families and 16 orders within the approximately 303 acres MU campus, demonstrating that a relatively small peri-urban academic landscape can support a taxonomically diverge avian assemblage (Savard et al. 2000). The occurrence of 37 species groups across a heterogeneous mosaic of dense tree canopy, scattered woodland areas, grasslands, wetlands, open fields and built-up areas suggests that the campus provides a range of habitats and ecological resources suitable for birds with diverse habitat and dietary requirements (Tews et al. 2004). These findings are particularly relevant in the context of Bodh Gaya, a region of considerable cultural and tourism importance and part of a landscape containing several natural and semi-natural habitats. The observed diversity also compares favourably with reports from other university campuses in Bihar and elsewhere in India. The 78 species recorded at MU were comparable to the 91 species documented at Nalanda University and the 81 species reported from the Central University of South Bihar (Yadav et al. 2024; Imran et al. 2025). Comparable levels of avian richness have also been reported from other Indian academic campuses (Chakdar et al. 2016; Sailo et al. 2019; Rathod and Bhaduri 2022; Kumar et al. 2024). Differences among campuses are likely to reflect variation in campus size, vegetation structure, surrounding land use, habitat connectivity, water availability, sampling duration, and methodological effort. Nevertheless, the consistently substantial bird richness reported from academic campuses reinforces their potential contribution to biodiversity conservation within human-dominated landscapes (Guthula et al. 2022).

Passeriformes constituted the most species-rich order, accounting for nearly half of the recorded species. The predominance of Passeriformes is consistent with their broad ecological radiation and ability to exploit structurally diverse habitats and a wide range of food resources (Tews et al. 2004; Zafar et al. 2026). The representation of numerous other orders, although at lower richness, further indicates that the campus is not utilized exclusively by a narrow group of urban-adapted birds but accommodates species with contrasting ecological requirements. The high representation of Muscicapidae at the family level may be associated with the availability of structurally complex vegetation and arthropod-rich foraging habitats, although such relationships needs to be further investigated (Basavarajappa et al. 2023). Overall, the taxonomic composition reflects the heterogeneous nature of the campus habitat and its capacity to support birds occupying multiple ecological niches.

The community level abundance pattern revealed an uneven distribution of individuals among species, with a relatively small number of species accounting for a substantial proportion of total abundance and a longer tail comprising less abundant species (Buckland et al. 2011). The steep initial decline in the rank-abundance curve is consistent with a community containing numerically dominant species alongside a large number of less abundant taxa. *Argya striata* (Jungle Babbler), *Acridotheres tristis* (Common Myna), *Spilopelia chinensis* (Spotted Dove), *Pycnonotus cafer* (Red-vented Bulbul), and *Cinnyris asiaticus* (Purple Sunbird) were among the most abundant species (Mahanta et al. 2025). Their prominence may reflect their ability to exploit a combination of natural, landscaped, and anthropogenic resources available within the campus (Verma and Murmu 2015). At the same time, the occurrence of numerous less abundant species indicates that the campus also provides opportunities for species with more specialized or less frequently encountered resource requirements. Thus, species richness and numerical dominance represent complementary aspects of the campus avian community, with high richness occurring alongside an uneven distribution of individuals.

The feeding-guild composition further demonstrated functional heterogeneity within the avian assemblage. Insectivores represented the largest guild, followed by omnivores, carnivores, frugivores, and granivores. The strong representation of insectivorous birds indicates that arthropod-based resources constitute an important component of the available foraging niche, while the presence of multiple additional feeding guilds suggests the availability of diverse food resources across the campus (Magura et al. 2010; Nyffeler et al 2018; Mahanta et al 2024). Insectivorous birds feed upon the terrestrial insects living near the ground level (Magura et al. 2010); aerial insects preferring the heights near the trees and shrubs; and the aquatic insects found near the water bodies at the campus. The frugivorous and granivorous species are supported by the fruiting trees and agricultural fields located in and around the study area. The substantial representation of omnivorous species may also reflect the ability of these birds to exploit both natural and human-associated food resources in a heterogeneous urban environment. The diversity of feeding guilds demonstrates that the campus supports birds occupying different trophic and resource-use niches. The observed pattern of avifaunal feeding guild appears to be closely associated with the heterogeneous habitat at the MU campus (Panda et al. 2021). The mosaic vegetation structure provides a broad range of residence and food resources to support a large assemblage rich diversity throughout the year.

Residential status revealed a predominantly resident avifauna, with 66 species classified as residents, compared with 11 winter migrants and one summer migrant. The strong predominance of resident species suggests that the campus provides sufficient habitat and resources to sustain a substantial component of its avian community throughout the year. The occurrence of winter migrants, however, demonstrates that the campus also provides seasonally suitable resources during periods of increased regional movement (Bisen et al. 2026). The much smaller representation of summer migrants is consistent with the generally greater prominence of winter movements in many parts of northern and eastern India. The presence of both resident and migratory species therefore emphasizes the dual role of the campus as a year-round habitat and a seasonal refuge within a rapidly urbanizing landscape (Ur Rahman et al. 2026). The presence of several species of aquatic birds during the monsoon season further indicates seasonal migration which is supported by formation of temporary water bodies during the rainy days.

The annual occurrence frequency analysis provided a complementary perspective on temporal consistency in their presence within MU campus. 51 out of 78 recorded species were of rare sighting. This predominance of a large proportion of rare sightings can be influenced by temporal occurrence pattern, habitat associations, seasonal factors, making MU campus as a dynamic green corridor for regional migration. A relatively small proportion of abundant (11.5%) and common (3.85%) occurring species suggests that a limited number of species were consistently sighted throughout the year. These regularly detected birds indicated that the campus provides a baseline of permanent resources to sustain their population irrespective of the season. This is consistent with the habitat mosaic of the MU campus which provides a diverse residential and foraging guild which can be used differently by different species (Panda et al. 2021; Tang et al. 2026).

Ordination analyses further indicated that seasonal changes did not result in complete replacement of the avian community. The concentration of many species around the central region of the Correspondence Analysis ordination suggests broad occurrence across seasons, whereas species positioned closer to individual seasonal points showed comparatively stronger seasonal associations. The relatively close positioning of the seasonal points indicates substantial similarity in community composition among seasons. Similarly, PCA showed that PC1 accounted for 96.84% of the total variance and PC2 explained a further 1.50%, together capturing 98.34% of the variation in seasonal abundance. The steep decline in the scree plot therefore indicates that most seasonal variation was concentrated along the first principal component (Legendre and Gallagher 2001; Yao et al. 2020). These findings suggest that seasonal changes in the campus avifauna were expressed primarily through differences in abundance and the occurrence of particular taxa rather than through complete restructuring of the community.

Seasonal Simpson’s dominance values remained low and varied within a narrow range (0.016), further supporting the absence of pronounced numerical dominance by a few species during any particular season. The lowest dominance during the post-monsoon period indicated relatively greater evenness, whereas the slightly higher value during winter suggested a modest increase in the contribution of some species. Such a pattern may be associated with seasonal movements, resource availability, or temporary aggregation of particular species. However, the small difference among seasons indicates that the overall distribution of individuals remained relatively even throughout year. Together with the CA and PCA results, these findings suggest that seasonal variation occurred within a broadly persistent community structure rather than through substantial seasonal replacement of species.

The conservation assessment provided an additional dimension to the ecological significance of the campus. Although 77 of the 78 recorded species were classified as Least Concern, one species, *Coracias benghalensis*, was categorized as Near Threatened. In addition, 19 recorded species showed declining global population trends. The predominance of Least Concern species should therefore not be interpreted as an absence of conservation concern, particularly where local populations may be influenced by habitat modification, declining resource availability, or broader landscape-level pressures. The occurrence of a Near Threatened species and multiple species with declining global trends highlights the value of maintaining mature vegetation, seasonal wetlands, open foraging areas, and other habitat elements that collectively sustain the campus avifauna (Chazdon et al. 2009; Tang et al. 2026).

The ecological value of MU campus appears to arise not from any single habitat type but from the interaction of multiple habitat elements within a relatively compact landscape. Mature trees provide nesting, roosting, and foraging opportunities; dense vegetation offers cover and structurally complex habitats; grasslands and open areas support ground-foraging species; buildings provide nesting and roosting sites for urban-associated birds; and seasonal water bodies provide additional resources for water-associated and migratory species. Such habitat heterogeneity can increase the range of available ecological niches and allow species with contrasting habitat and resource requirements to coexist within the campus (Cramer and Willig 2005; Tang et al. 2026).

Overall, these findings demonstrate that the peri-urban MU campus supports a diverse and relatively persistent avian assemblage within a rapidly urbanizing landscape, highlighting its ecological and conservation significance in the Gangetic plains of Bihar. Its predominantly resident avian assemblage, representation of multiple feeding guilds, occurrence of seasonal migrants, persistence of a core species component, and presence of a Near Threatened species collectively emphasize the importance of retaining and managing its heterogeneous habitats. The enhancement of native vegetation, mature trees, heterogeneous habitats, and seasonal water bodies, together with the minimization of unnecessary anthropogenic disturbance, can help sustain avian diversity and functional heterogeneity in rapidly transforming urban environments. Integrating biodiversity-sensitive habitat management and ecological restoration into campus planning may further strengthen the role of university landscapes as refugia for wildlife and providers of essential ecosystem functions.

## Statements & Declarations

## Author Contributions

All authors contributed to the conception and design of the study and to data analysis. Data collection was performed by Manisha Kumari. The first draft of the manuscript was written by Manisha Kumari and all authors contributed to the subsequent versions of the manuscript by providing their comments and revision. Kumari Aditi and Partha Pratim Das supervised the research work. All authors critically reviewed and approved the final version of the manuscript.

## Data Availability

The data supporting the findings of this study are available from the corresponding authors upon reasonable request.

## Consent to Participate

Consent to Participate declaration: not applicable.

## Ethics Approval

Not applicable. This study involved non-invasive field observations of birds and did not involve the capture, handling, marking, experimentation, or any other invasive procedures on animals.

## Acknowledgements

The authors sincerely acknowledge the Post Graduate Department of Zoology, Magadh University, Bodh Gaya for providing the necessary facilities and institutional support for conducting field surveys and completing the present study. Manisha Kumari acknowledges the Council of Scientific and Industrial Research – University Grants Commission (CSIR-UGC) for providing Junior Research Fellowship (JRF) (E-certificate no. 24J/03/01082) to carry out this research work.

